# Spatial mapping of cell types by integration of transcriptomics data

**DOI:** 10.1101/2019.12.13.874495

**Authors:** Alma Andersson, Joseph Bergenstråhle, Michaela Asp, Ludvig Bergenstråhle, Aleksandra Jurek, José Fernández Navarro, Joakim Lundeberg

**Affiliations:** Science for Life Laboratory, KTH Royal Institute of Technology, School of Engineering Sciences in Chemistry, Biotechnology and Health, Department of Gene Technology, 171 65 Solna, Sweden

## Abstract

Spatial transcriptomics and single cell RNA-sequencing offer complementary insights into the transcriptional expression landscape. We here present a probabilistic method that integrates data from both techniques, leveraging their respective strengths in such a way that we are able to spatially map cell types to a tissue. The method is applied to several different types of tissue where the spatial cell type topographies are successfully delineated.

## 1 Main

Techniques for spatial transcriptomics have advanced to a state where the entire transcriptome now can be spatially resolved, however methods providing an exhaustive portrait of the expression with deep coverage do not yet guarantee resolution at the single cell level. [1–3] Thus transcripts captured at a given position may stem from a heterogeneous set of cells, not all necessarily of the same type. Hence the observed expression profile at any location can be considered a mixture of transcripts originating from multiple distinct sources. Implicitly this means that even though the transcriptional landscape can be thoroughly charted, the biological identity and spatial distribution of the cells generating this remains largely unknown.

Spatial transcriptomics techniques face a dilemma of knowing the location of transcripts but not which cell that produced them, while the opposite is true for data retrieved from single cell RNA-sequencing experiments; where each transcript is associated to an individual cell but information regarding their position within the tissue is lost. Given this set of complementary strengths and weaknesses, the idea of combining data from the two techniques to delineate the spatial topography of cell type populations is compelling.

The method we present integrates single cell and spatial transcriptomics data, the latter originating from the technique presented by Ståhl et.al and referred to as ST, allowing cell types to be spatially mapped onto a tissue. [3,4] More explicitly, the proportion of each cell type at a capture location (hereafter; spot) is determined by a probabilistic model informed by both types of data. In short, we first define expression profiles that are characteristic of the cell types and then find the combination that best reconstruct the observed spatial data, eliminating any need for interpretation of abstract concepts like factors or clusters. [5] Although methods to deconvolve (bulk) RNA-seq data have already been presented and could theoretically be applied to ST data, these tend to exhibit certain limitations such as: only certain cell types can be assessed, manual curation of data is required to form representative cell type “signatures”, no clear theoretical foundation behind the method exists or (as mentioned above) the components into which the data is decomposed lack a clear biological interpretation. [5–8]

Raw count matrices for the single cell and ST data together with annotations of the former are the only three items necessary to conduct our analysis, no normalization or other transformations are applied. Any combination of single cell and ST data sets of similar composition can be used, without the need for them to be paired (i.e. from the same tissue specimen), allowing publicly available resources to be fully utilized. The method rests on the primary assumption that both ST and single cell data follow a negative binomial distribution, commonly used to model expression count data. [5, 9] Technical bias is taken as independent of cell type, and the types’ underlying expression profiles are seen as inherent biological properties unaffected by the method of choice to study them.

Two distinct steps constitute the actual implementation; first parameters of the negative binomial distribution are estimated from the single cell data, for all genes within each cell type. Equivalent parameters for a distribution describing the expression from a mixture of these types, like that in a spot, can be obtained by a weighted combination of the single cell parameters. In the second step such weights are estimated with a condition that the resulting distribution should provide the best possible explanation of the provided ST data. Cell type proportions are obtained by normalizing the weights to make them sum to unity (Methods). Partitioning the process into two distinct steps has the advantage that once single cell parameters have been estimated, they can be applied to any ST data set of choice without the need to be re-estimated. Fig. 1 displays a schematic overview of the workflow upon using the implemented method.

**Figure 1:**
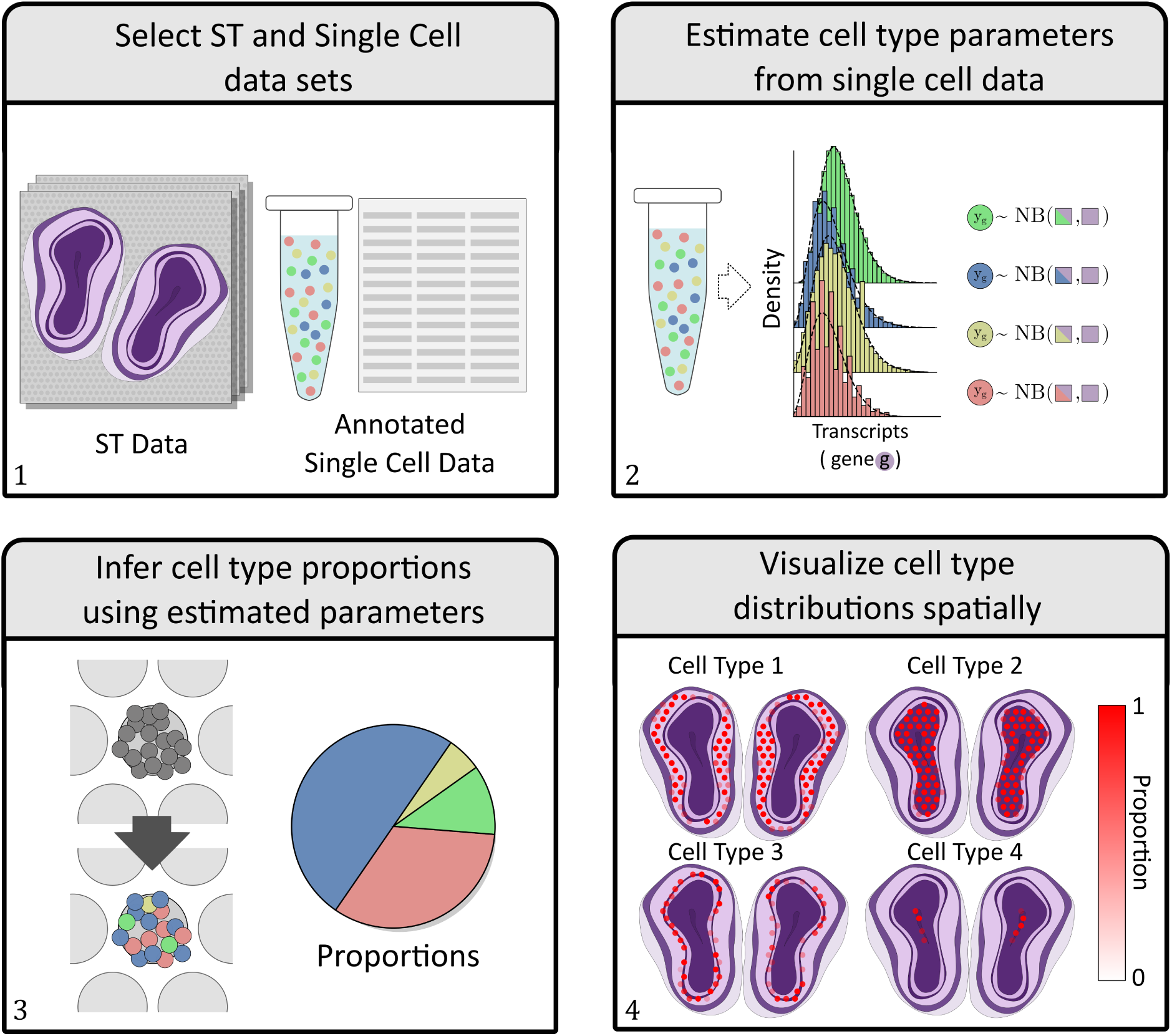
Schematic Overview of method workflow. (1) Annotated single cell data and a set of ST data with similar cell type content are selected for analysis. (2) Parameters of the negative binomial distributions characterizing the expression are estimated from the single cell data – with the first parameter (the rate) being conditioned on both gene and cell type whilst the second (the success probability) is only conditioned on gene (Methods). (3) Estimated parameters are used to infer cell type proportions in each spot. (4) The spatial organization of cell types are visualized by letting the opacity of each spot’s face color represent the proportion values.

In order to show the utility of the method we apply it to two different tissues: human developmental heart (6.5 post conceptional weeks, PCW) and mouse brain. Furthermore, we only use ST and single cell sets derived from disparate sources to illustrate how paired data is not required to render factual results. See Methods for complete specifications.

We consider the developmental heart and mouse brain tissues as good candidates to evaluate the method. The developmental heart’s anatomy has been thoroughly explored and previous studies provide insights into the expected location of certain cell types. As for the mouse brain, it has also been extensively studied; resulting in plenty of resources describing its anatomical and molecular properties, one of them being the Allen Brain Atlas (ABA). [10] By combining information of known cell type marker genes with the available ISH (In Situ Hybridization) data in ABA, the expected spatial distribution of these types can be deduced and used as a reference to compare our results with. Figure 2 displays a subset of the results obtained upon mapping the single cell data onto the mouse brain ST data sets (complete analysis in Supplementary Section 1.2.2). Each spot is represented by a circle where the alpha-level of the facecolor indicates how abundant a certain cell type is at the given location, i.e. the higher the opacity, the higher the estimated proportion of the studied cell type (Methods). As shown in Figure 2A, single cell clusters can be mapped onto the tissue, informing us of what spatial patterns they exhibit and how these clusters physically relate to each other — the spatial context may also aid in assigning more distinct and descriptive identities to the clusters.

**Figure 2:**
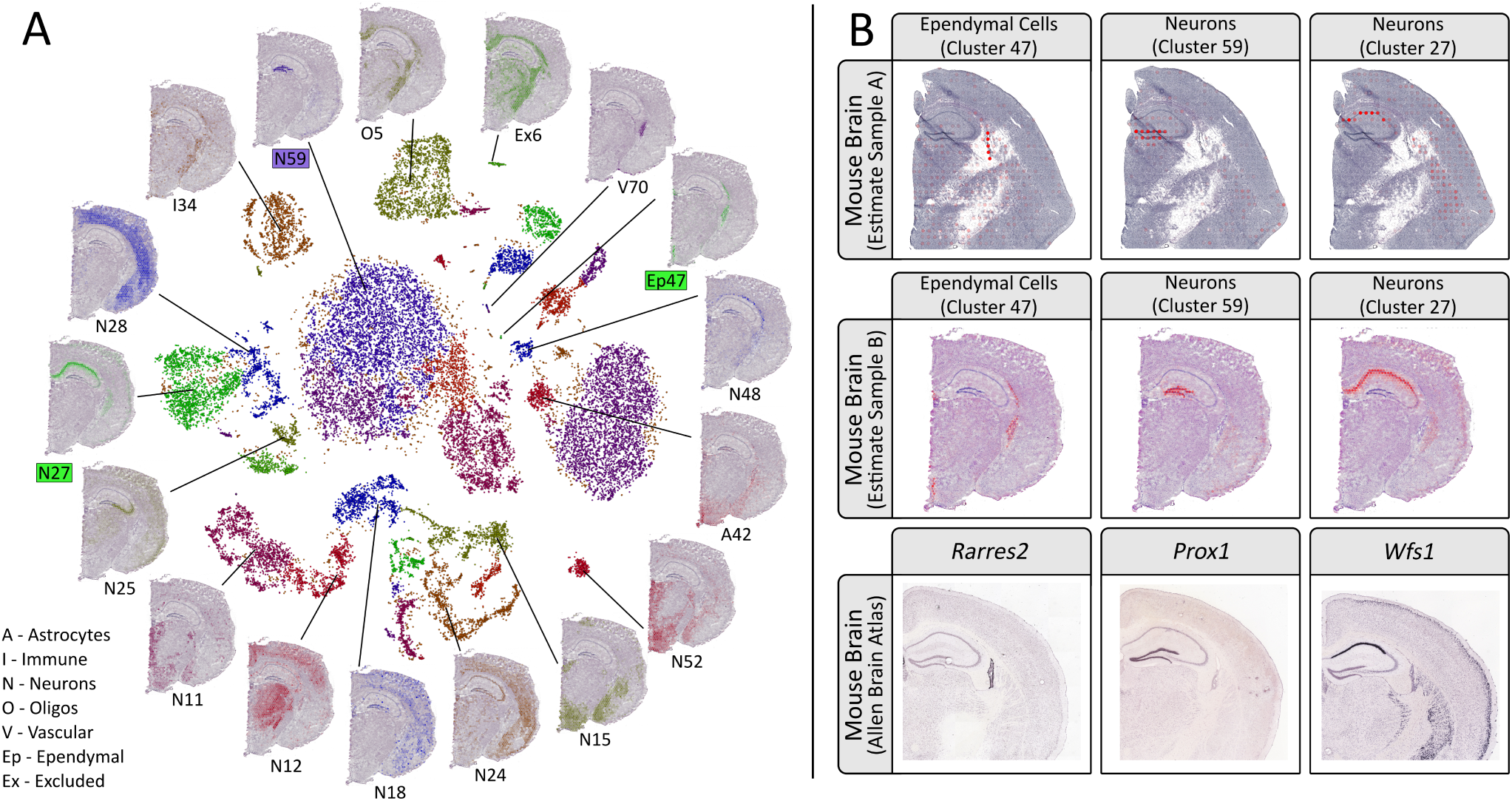
**A)** Visualization of the single cell hippocampus data by using its gt-SNE embedding (inner region), with spatial proportion estimates of several clusters overlaid on the H&E-image (outer region) of sample mb-B (10x Visium array, 55 micron spots). The cluster labels are derived from the original single cell data set (Methods). [11,12] **B)** Estimated proportions for three of the 56 clusters, (here taken as cell types), defined in the mouse brain single cell data set. Two different sections are used mb-A (ST array, 100 micron spots) and mb-B, to illustrate the consistency between different array resolutions. Marker gene expression patterns are obtained by ISH are found in the bottom row, taken from the Allen Brain Atlas. *Rarres2* is a marker gene of Ependymal cells, *Prox1* for Dentate Granule Neurons and *Wfs1* for Pyramidal Neurons (the latter two both being subtypes of neurons).

When assessing our results for the mouse brain, *Rarres2* is taken as a marker gene for Ependymal cells (cluster 47), *Prox1* for Dentate Granule Neurons (cluster 59) and *Wfs1* for Pyramidal Neurons (cluster 27); only broad classes like “Neurons” are provided in the single cell data annotations, but observing their spatial arrangement enables us to assign more granular types of these classes to the clusters. [13–15] It’s evident how the estimated proportions agree with the signals observed in the ISH experiments, confirming the proposed locations of these cell types. There is a high degree of consistency of the mapping between the different sections that are analyzed, speaking in favor of the method’s robustness. In addition to coinciding with marker gene expression, the suggested spatial organization is further supported by already established knowledge regarding these types. Ependymal cells line the ventricular system, forming an epithelial sheet known as the ependyma, thus observing strong signals for this cell type in the lateral ventricular region is affirmative. [16] Dentate Granule Neurons reside within the dentate gyrus, as implied by their name, a feature that our mapping manages to reproduce. [17] Pyramidal Neurons belong to the broad class of excitatory neurons and populate regions such as the amygdala, cerebral cortex and parts of Ammon’s horn in the hippocampus, again in line with our results. [18] The usefulness of our method might be argued in a scenario where the marker gene(s) of types are known, since in theory expression levels could simply be visualized and used to infer the types’ presence. However, due to the sparsity and variance in ST data this single gene approach does not always manage to recreate the patterns observed in ABA (see Supplementary section 1.2.2, Supplementary Figure 16,17 and 18), attesting to how using the full expression profiles of cell types is preferable to relying on a few genes when working with this kind of data.

In the developmental heart (Supplementary Section 1.2.1) we observe how ventricular and atrial cardiomyocytes have the highest proportion values in the ventricular body respectively the atria. From the H&E images (Hematoxylin and Eosin), blood cells are visible within the hollow cavities, the same areas as they are mainly estimated to reside within. Smooth muscle cells are almost exclusively mapped to the outflow tract, again, in concordance with their expected location. [19] Epicardial cells form a thin outer layer of the heart known as the epicardium, and this type is mainly assigned high proportion values in spots covering the edges of the heart. [20] Epicardium-derived cells arrange themselves adjacent to the epicardial cells on the inner side of the heart in a somewhat thicker layer than the epicardium, and they are also known to be present in the outflow tract during its formation, a pattern recapitulated by our results. [21]

Finally we generated synthetic ST data from single cell data (Methods), providing us with a “ground truth” for the proportion values. The synthetic data enabled a comparison with two other recently published methods designed to deconvolve bulk data with the help of single cell data (DWLS and deconvseq); where our implementation performed better than both of the other methods (Supplementary section 1.2.3). [8, 22]

Once the proportions have been estimated, subsequent analysis supplementary to visualization can be conducted. To give one example, by looking at the spatial correlation between cell types (Pearson correlation on a spot basis, Methods) we can investigate which cell types that tend to co-localize together and potentially interact with each other. This is a complementary approach to that of using receptor-ligand pairs to assess cell type interactions in a sample, without the need for curation of lists describing receptors, ligands and their interactions. [23]

To summarize, we present both a method and implementation to map cell types found in single cell data spatially onto a tissue. Our implementation is released as a python package named *Stereoscope* available at *github.com/almaan/stereoscope*. The procedure is seamless and compatible with any two data sets, does not require any processing of the data and, albeit we focus on ST data here, the method is applicable to any method where the observed transcriptomics data can be considered a mixture of contributions from multiple individual cells added together in a linear manner.

## 2 Methods

### 2.1 Code Availability

The method is released as a tool named *Stereoscope* available at: https://github.com/almaan/stereoscope.

Documentation for *Stereoscope*, a tutorial, scripts used for visualization and further analyses are also found within the repository. In the tutorial we provide walk-throughs to reproduce some of the analyses presented in this paper, from the very first step of downloading data to visualizing the results.

### 2.2 Model

The following notations will be used upon describing the model

- *G* – the set of all genes
- *S* – the set of all spots
- *Z* – the set of all cell types
- *C*_*s*_ – the set of all cells contributing to spot *s*
- *n*_*sz*_ – number of cells from cell type *z* at spot *s*
- *x*_*sg*_ – counts of gene *g* at spot *s*
- *x*_*sgc*_ – counts of gene *g* at spot *s* from cell *c*
- *z*_*c*_ – cell type of cell *c*
- *α*_*s*_ – scaling factor at spot *s*
- *β*_*g*_ – technique based gene bias for gene *g*
- *r*_*gz*_ – rate parameter for cell type *z* and gene *g*
- *p*_*g*_ – success probability parameter for gene *g*
- | · | – cardinality of a given set
- ***a***- vector notation

Transcripts of a given gene (*g*) within a single cell (*c*) are taken as negative binomially distributed – with the rate 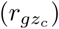 being conditioned on a cell’s type (*z*_*c*_) and gene *g*, whilst the success probability is only dependent on the gene in question (a common postulation). [9, 24] To account for certain technical biases, we also include a cell specific scaling factor *s*_*c*_, taken as the reciprocal of each cell’s library size. Thus we have

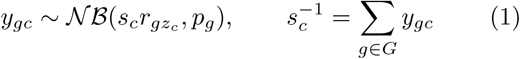

Values for the cell type specific parameters are then obtained by finding the MLE (maximum likelihood estimates), given the provided single cell data. In the implementation this is achieved by taking the negative log-likelihood as an objective function to be minimized w.r.t the parameters. PyTorch is used for the optimization. [25]

In ST data, the observable transcripts (*x*_*sgc*_) of a given gene (*g*) from a cell (*c*) contributing to a specific spot (*s*) are also taken as negative binomially distributed, with the same conditioning as for the single cell data. We assume that the efficiency by which certain genes are captured differs between the two techniques (ST and single cell RNA-seq), what would be referred to as technique based bias, and thus introduce a variable (*β*_*g*_) to correct for this. A scaling factor (*α*_*s*_) for each spot is also included to account for technical variation between the spots. The distribution used to model the ST data thus takes the form

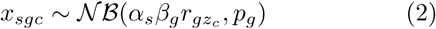

The total number of transcripts (*x*_*sg*_) for a certain gene (*g*) at each spot (*s*) is simply the sum of observed transcripts from each cell (*c*) contributing to that spot, that is

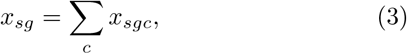

With a shared second parameter (*p*_*g*_) between all types (*z*), the first parameter exhibit an additive property and the total number of transcripts can be taken as negative binomially distributed as well

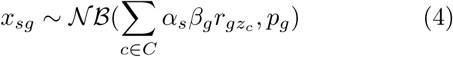

By introducing a quantity coefficient *n*_*sz*_ representing the number of cells from a certain type (*z*) present at spot *s*, a change of index from *cells* to *types* is possible

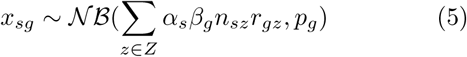

We then bundle the spot specific parameters together in a scaled quantity coefficient (*v*_*sz*_)

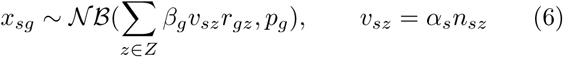

Using vector notation this expression can be rewritten as

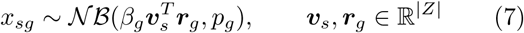

To account for asymmetric data sets (where the cell types in ST and single cell data do not overlap perfectly) and noise we also include a form of “dummy” cell type, with gene specific rates (*ϵ*_*g*_) and a scaled quantity coefficient *γ*_*s*_.

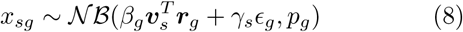

If we define *w*_*sz*_ as the normalized scaled quantity coefficients, excluding the noise capturing dummy cell type, that is

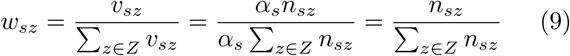

this results in an expression which can be recognized as the proportion of each cell type within a given spot.

To avoid promiscuous assignment of explanatory power to the dummy cell type, we place a standard normal prior on all of its rates, i.e.

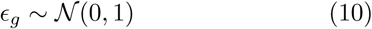

Cell type proportions (*w*_*sz*_) are then taken as the MAP (maximum a posteriori) estimate of the distribution in Eq. 8 using the prior in Eq. 10, given the observed ST data. Uniform priors are assigned to all other variables. More precisely this is implemented by minimizing the negative logarithm of the posterior w.r.t. to the scaled quantities ({***v***_*s*_}_*s*∈*S*_), the gene specific bias (***β***) and parameters related to the dummy cell type (***γ*** respectively ***ϵ***). Similar to the procedure for single cell data, the optimization is performed using PyTorch.

### 2.3 Processing Data

Here we give a description of how the data was processed; note that our “starting material” are raw count matrices of single cell and ST data, in formats [*cells*]x[*genes*] respectively [*spots*]x[*genes*] where the single cell set has some form of meta-data containing type annotations. For exact details regarding how these count matrices were obtained from the raw sequencing data, we refer to their original publications.

#### 2.3.1 Human Developmental Heart

The complete single cell data set provided in the paper *“A spatiotemporal organ-wide gene expression and cell atlas of the developing human heart”*, was used to estimate the type parameters hence resulting in a usage of 3717 cells distributed over 15 clusters. [20] Only the top 5000 highest expressed genes were used in the analysis. For the exact composition of the single cell data set, see Supplementary section 1.1.1.

ST data was taken from the same publication as the single cell data, using the 8 sections from PCW 6.5. Only those spots under the tissue were used. From the 5000 genes selected in the single cell data, the intersection of these and the complete set of genes found in the ST data was used.

#### 2.3.2 Mouse Brain

The single cell data set was downloaded from mousebrain.org, where we used the data containing cells originating from Hippocampal tissue. [11] We first joined the “Class” and “Cluster” identifiers for each cell to form type labels. A subset of 8449 cells were sampled from the 29519 cells found within the set. This subset was assembled by specifying both a global lower (*l*) and upper (*u*) bound for the number of cells to be included from each type, and then applying the procedure given in Eq 11 (*n*_*z*_ representing the total number of cells from type *z*). We use an upper bound to reduce run time.

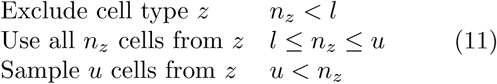

The lower and upper bounds were set to 25 respectively 250 cells, giving the subset a total of 56 clusters. Only the top 5000 highest expressed genes were used in the analysis. See Supplementary section 1.1.2 for a more detailed description of the set composition.

From the ST data, only those spots under the tissue were used. Three sections (mb-A, mb-*α* and mb-B) are used in the analysis. From the 5000 genes selected in the single cell data, the intersection of these and the complete set of genes found in the ST data were used. mb-A and mb-*α* were analyzed together whilst mb-B was analyzed separately.

#### 2.3.3 ISH Images

ISH images were downloaded from the Allen Brain Atlas. No modifications except for cropping were applied. References for the used images are:

- *Rarres2* [26]
- *Prox1* [27]
- *Wfs1* [28]

### 2.4 Synthetic ST Data and Comparison

#### 2.4.1 Method

To allow for comparison of performance between methods we devised a method for generation of synthetic data. We refrain from using any approach based on a negative binomial model, as this potentially could favour our model in an unfair way. Thus we decided to rather use a “semi-synthetic’ approach not based on a negative binomial model, where we use single cell data to produce synthetic ST data. The procedure is described in Algorithm 1.

##### Algorithm 1 Synthetic data generation

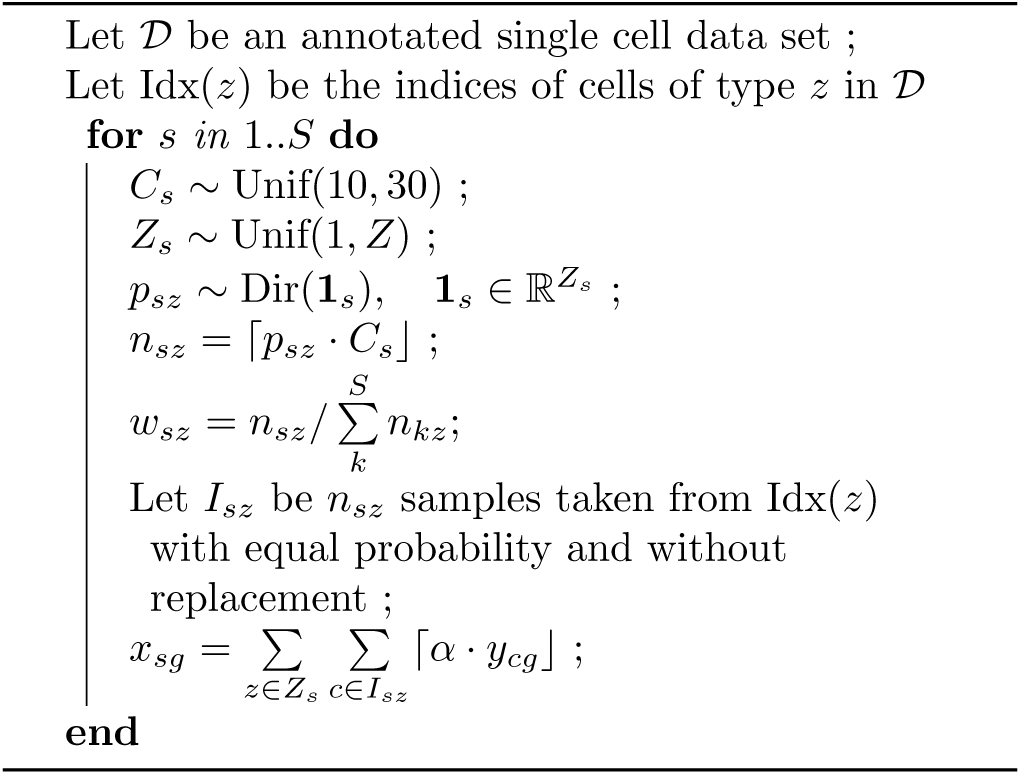

Meaning that for every spot (*s*) we first sample the number of cells (*C*_*s*_) contributing to this, and the number of types (*Z*_*s*_) which these cell may belong to. We then draw unadjusted proportions from the probability simplex using a Dirichlet distribution (concentration set to 1 for all present types). The actual number of cells from each type (*n*_*sz*_) are then set to the nearest integer number for the corresponding proportion of cells in the spot. The adjusted proportions (*w*_*sz*_) are given as the actual proportion based on the number of cells after the nearest integer rounding. From each cell type (*z*) we then sample (without replacement) indices (*I*_*sz*_) from cells in the single cell data set that are labeled as this type (Idx(*z*)). To generate the expression value for each gene (*x*_*sg*_) we sum the the nearest integer approximation of the product between the single cell expression values (*y*_*sg*_) and a scaling factor (*α*_*s*_), a constant specified by the user, over all selected types and the sampled indices. By applying this procedure one obtains an ST data set where the “ground truth” regarding the proportions is known (the adjusted proportions).

#### 2.4.2 Generated Set

The single cell data set we used was that of hippocampus taken from mousebrain.org (same as for the mouse brain analysis), where the annotations used were those labels given as “Subclass”. We first subsampled the set according to the procedure described above (using 60 as lower respectively 500 as upper bound). The subsampled set was then split into two equally sized and mutually exclusive sets, i.e. sharing no cells. We refer to these as the *generation* and *validation* set. A synthetic ST data set was then generated according to the procedure outlined in Algorithm 1 using the generation set as input. The resulting ST data set contained 1000 spots and 500 genes (the top 500 highest expressed). The purpose of the validation set is to be used as the single cell data provided together with the ST data as input to respective method.

#### 2.4.3 Comparison and Evaluation

To compare the performance between methods, we provided each of them with the validation single cell data set and the generated synthetic ST data to obtain proportion estimates for each spot. For each method we then computed the RMSE (Eq.12) between the estimated proportions (***w***) and the ground truth 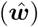.

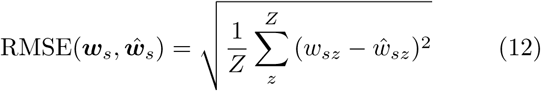

Being interested in whether our method performed better than the others, we conducted a one sided Wilcoxon signed-rank test, to see whether the difference between RMSE values in each spot are asymmetrically distributed around zero – in favor of our method. This is done using the scipy implementation of the Wilcoxon signed-rank test. [29] Two recently published methods were selected for comparison: *DWLS* and *deconvSeq*. [8, 22] Slight modifications had to be made to the code in DWLS, though these changes did however not concern the actual proportion estimation. All code used throughout the comparison, including wrappers for the methods when applying them to ST data, are found in the github repository. The aforementioned modifications are accounted for in more detail at said repository. To put the RMSE values into context, we compute the RMSE between probabilities drawn from a Dirichlet distribution (all concentration values set to 1) for an equal number of spots as in the analyzed data sets. By repeating this for a select number of times, we obtain a “null-distribution” of RMSE values, to compare the other RMSE-distributions to.

### 2.5 Reproducing the analysis

Below, we describe specific details for the analysis of each pair of datasets, allowing the results to be properly reproduced

|  | Developmental<br>Heart | Mouse<br>Brain | Breast<br>Cancer |
| --- | --- | --- | --- |
| SC Epochs | 50000 | 50000 | 50000 |
| ST Epochs | 50000 | 50000 | 50000 |
| Learning Rate | 0.01 | 0.01 | 0.01 |
| Top N Genes | 5000 | 5000 | 5000 |

## 3 Software

Code is written in Python 3.7, the core functions rely on the following libraries (and built with versions): numpy 1.17.4, torch 1.3.1 and pandas 0.25.3. Additional libraries for tasks such as parsing and logging are used for the CLI application and visualization; the entire list is given at the github repository and included in the installation file.

## 4 Visualization and Further Analysis

All results for the proportion estimates are visualized by the same procedures, modules and scripts for this are provided at the github page.

### Proportions – separate visualization

Upon visualizing the proportion of a single type within a given spot the opacity for the red face color corresponds to the estimated proportion. Proportion values are scaled within each section and cell type, to emphasize the spatial patterns of each cell type within the tissue. Hence for the set of proportion values within each cell type and section, all elements are divided by the largest element found within said set, adjusting the range of the values to the unit interval. No threshold or adjustment of the values is applied. The spots coordinates are transformed to pixel coordinates and overlaid on the H&E image.

### Proportions – joint visualization

We use a previous propsed method for visualization of higher-dimensional spatial data. [30] This enables a joint visualization of the cell type distributions to be produced, where regions of similar colors share similar compositions of cell types. The procedure consists of two steps: (1) an embedding of high-dimensional data points to a 3-dimensional manifold (*f*: ℝ^|*Z*|^ 1↦ ℳ^3^) and (2) a transformation *g*: ℳ^3^ ↦ [0, 1]^3^ corresponding to a mapping of values into the unit cube; Eq. 13 gives a more explicit description.

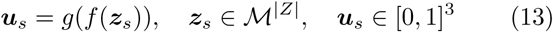

We choose *f* to preserve some of the data’s internal structure, the resulting three dimensional vector ***u***_*s*_ is then used as RGB values, using the first element as the red channel value, the second as green and third as blue. Examples of such “structure preseving” mappings are popular dimensionality reduction techniques such as tSNE and UMAP. [31, 32] Due to our choice of *f*, colors are indicative of cell type composition. It’s important to note how *a color* does *not* necessarily correspond to a single cell type. Whenever any of the terms “compressed visualization” or “joint visualization” are used, this is the type of visualization we refer to.

### Hippocampus Single Cell – Cluster Visualization

To generate the image presented in Figure 2A, we used the coordinates obtained upon embedding the data within a 2-dimensional manifold using gt-SNE. [11] These coordinates were provided in the single cell data loom-file, as attributes named “_X” and “_Y” respectively, and hence were not generated by us. The cluster indices are those obtained upon joining the “Class” and “Cluster”identifiers for each cell. Clusters excluded from the proportion estimate analysis are not visualized in the gt-SNE plot. The proportion estimates are those obtained upon analyzing the mb-B section together with the single cell data set as described in Methods 2.3.2.

### 4.1 Correlation of cell type proportions

By computing the Pearson correlation (See Eq 14) between each pair of cell types, treating each spot as a distinct data point, one obtains information regarding which cell types that share a similar spatial distribution.

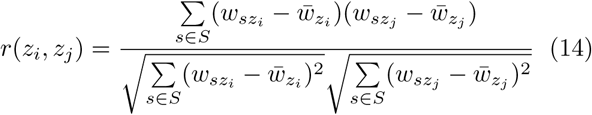

In Eq. 14 *z*_*i*_ represents cell type *i*, the bar indicates the arithmetic mean, and *S* is the set of spots in the studied data set. Where *s* represents a specific spot and *w*_*sz*_ the proportion of cell type *z* in said spot.

## 4.2 Acknowledgements

We want to thank Vilhelm Bohr, Tyler Demarest and Deborah Croteau for providing us samples of the Mouse Brain. Additional thanks are also given to The Knut and Alice Wallenberg (KAW) Foundation, the Thon Foundation, EU JPND INSTALZ, Foundation for Strategic Research (SSF), Science for Life Laboratory and the Royal Institute of Technology (KTH) who enabled this work to be produced.

## 4.3 Contributions

A.A. formulated the model, implemented it in code and wrote the manuscript. J.F.N. and A.J. contributed with the mouse brain ST-data. M.A. provided early access to the developmental heart single cell and ST data, aided in assessing the results obtained for the same tissue and commented on the manuscript. J.B., J.F.N. and L.B. gave comments on the method and manuscript. J.L. supervised the project, commented on the manuscript and provided computational resources.

