## Supplementary Information for "Spatial mapping of cell types by integration of transcriptomics data"

### 1 Supplementary

#### 1.1 Data

The following section contains additional details about the data sets used; such as accession to the data and the exact composition of cell types within each single cell data set used.

##### 1.1.1 Developmental Heart

Both single cell and ST data for the analysis of the developmental heart are taken from the publication "*A spatiotemporal organ-wide gene expression and cell atlas of the developing human heart.*". The complete set of single cells was used, whilst only the 8 ST sections from PCW 6.5 were used. Supplementary Table 1 gives the specifics of the single cell data set.

| | Cell Type | Number of Cells |
|---|---|---|
| **1** | Atrial_cardiomyocytes | 152 |
| **2** | Capillary_endothelium | 662 |
| **3** | Cardiac_neural_crest_cells_Schwann_progenitor_... | 75 |
| **4** | Endothelium_pericytes_adventitia | 127 |
| **5** | Epicardial_cells | 128 |
| **6** | Epicardium-derived_cells | 392 |
| **7** | Erythrocytes_11 | 113 |
| **8** | Erythrocytes_6 | 186 |
| **9** | Fibroblast-like_cardiac_skeleton_connective_ti... | 463 |
| **10** | Fibroblast-like_larger_larger_vascular_develop... | 150 |
| **11** | Fibroblast-like_smaller_vascular_development | 337 |
| **12** | Immune_cells | 76 |
| **13** | Myoz2-enriched_cardiomyocytes | 97 |
| **14** | Smooth_muscle_cells_fibroblast-like | 263 |
| **15** | Ventricular_cardiomyocytes | 496 |
| | **Total** | **3717** |

Supplementary Table 1: Composition of the developmental heart single cell dataset.

##### 1.1.2 Mouse Brain

Single cell data was downloaded from *mousebrain.org*, where data for the Hippocampus was provided as a loom-file containing a total of 29519 cells, in contrast to the 18147 listed at the download page. As stated in the methods section the labels given as "ClusterID" and "Class" were joined together toin order to define cell types. Applying the subsampling scheme described in Methods, a dataset consisting of 8449 individual cells was assembled, for which the exact composition is given in Supplementary Table 2.

| | Cell Type | Number of Cells |
|---|---|---|
| **1** | Astrocytes_14 | 250 |
| **2** | Astrocytes_40 | 250 |
| **3** | Astrocytes_41 | 31 |
| **4** | Astrocytes_42 | 250 |
| **5** | Astrocytes_44 | 41 |
| **6** | Blood_73 | 46 |
| **7** | Ependymal_47 | 27 |
| **8** | Excluded_30 | 30 |
| **9** | Excluded_38 | 25 |
| **10** | Excluded_44 | 56 |
| **11** | Excluded_6 | 87 |
| **12** | Immune_14 | 63 |
| **13** | Immune_32 | 131 |
| **14** | Immune_34 | 250 |
| **15** | Immune_35 | 38 |
| **16** | Neurons_10 | 35 |
| **17** | Neurons_11 | 250 |
| **18** | Neurons_12 | 250 |
| **19** | Neurons_14 | 250 |

| | Type | Number of Cells |
|---|---|---|
| **20** | Neurons_15 | 250 |
| **21** | Neurons_16 | 52 |
| **22** | Neurons_17 | 250 |
| **23** | Neurons_18 | 248 |
| **24** | Neurons_19 | 55 |
| **25** | Neurons_20 | 38 |
| **26** | Neurons_21 | 250 |
| **27** | Neurons_22 | 27 |
| **28** | Neurons_23 | 249 |
| **29** | Neurons_24 | 250 |
| **30** | Neurons_25 | 250 |
| **31** | Neurons_26 | 250 |
| **32** | Neurons_27 | 250 |
| **33** | Neurons_28 | 250 |
| **34** | Neurons_30 | 26 |
| **35** | Neurons_48 | 183 |
| **36** | Neurons_49 | 25 |
| **37** | Neurons_51 | 250 |
| **38** | Neurons_52 | 241 |

| | Type | Number of Cells |
|---|---|---|
| **39** | Neurons_54 | 30 |
| **40** | Neurons_58 | 48 |
| **41** | Neurons_59 | 250 |
| **42** | Neurons_60 | 250 |
| **43** | Neurons_61 | 106 |
| **44** | Neurons_62 | 42 |
| **45** | Neurons_63 | 250 |
| **46** | Oligos_0 | 250 |
| **47** | Oligos_1 | 158 |
| **48** | Oligos_14 | 101 |
| **49** | Oligos_5 | 250 |
| **50** | Oligos_53 | 250 |
| **51** | Vascular_14 | 75 |
| **52** | Vascular_46 | 61 |
| **53** | Vascular_67 | 250 |
| **54** | Vascular_68 | 250 |
| **55** | Vascular_69 | 41 |
| **56** | Vascular_70 | 33 |
| | **Total** | **8449** |

Supplementary Table 2: Composition of the Mouse Brain data set.

We analyzed two 100 micron array ST sections (mb-A and mb-$\alpha$), data can be accessed at the github page, excluding all spots that did not cover the tissue. We downloaded Visium (55 micron array) data from the website of *10x Genomics*™, listed under "Support", "Spatial Gene Expression" and "Datasets", selecting the set listed as "Mouse Brain Section (Coronal)".[1] For the Visium data we only selected spots under the tissue.

###### 1.1.3 Synthetic Data

**Single Cell**
To generate synthetic single cell data, the data set originating from Hippocampal tissue was downloaded from mousebrain.org (the same set as used for analysis of the mouse brain) where the "Subclass" annotations were used as cell type identifiers. A generation and validation set of equal compositions (in terms of number of cells from each cell type) were generated, exact structure given in Supplementary Table 3.

| | Cell Type | Number of Cells |
|---|---|---|
| **1** | Astrocyte | 250 |
| **2** | Astrocyte,Neurons | 58 |
| **3** | Astrocyte,Oligos | 47 |
| **4** | Blood | 40 |
| **5** | Immune | 250 |
| **6** | Neurons | 250 |
| **7** | Neurons,Cycling | 94 |
| **8** | Neurons,Oligos | 33 |
| **9** | Oligos | 250 |
| **10** | Vascular | 250 |
| | **Total** | **1522** |

Supplementary Table 3: Composition of the generation and validation single cell data sets (the two share identical compositions).

**Spatial Data**
A total of 1000 spots were synthesized according to the procedure described in the Methods section. Only data from the top 500 highest expressed genes in the generation data set were used, hence a $1000 \times 500$ count matrix was generated. All synthetic data sets and the code used for the "synthesis" is available in the github repository of this paper, where also a tutorial to reproduce the results is presented.

[1] Link : https://support.10xgenomics.com/spatial-gene-expression/datasets/1.0.0/V1_Adult_Mouse_Brain).

#### 1.2 Results

##### 1.2.1 Developmental Heart

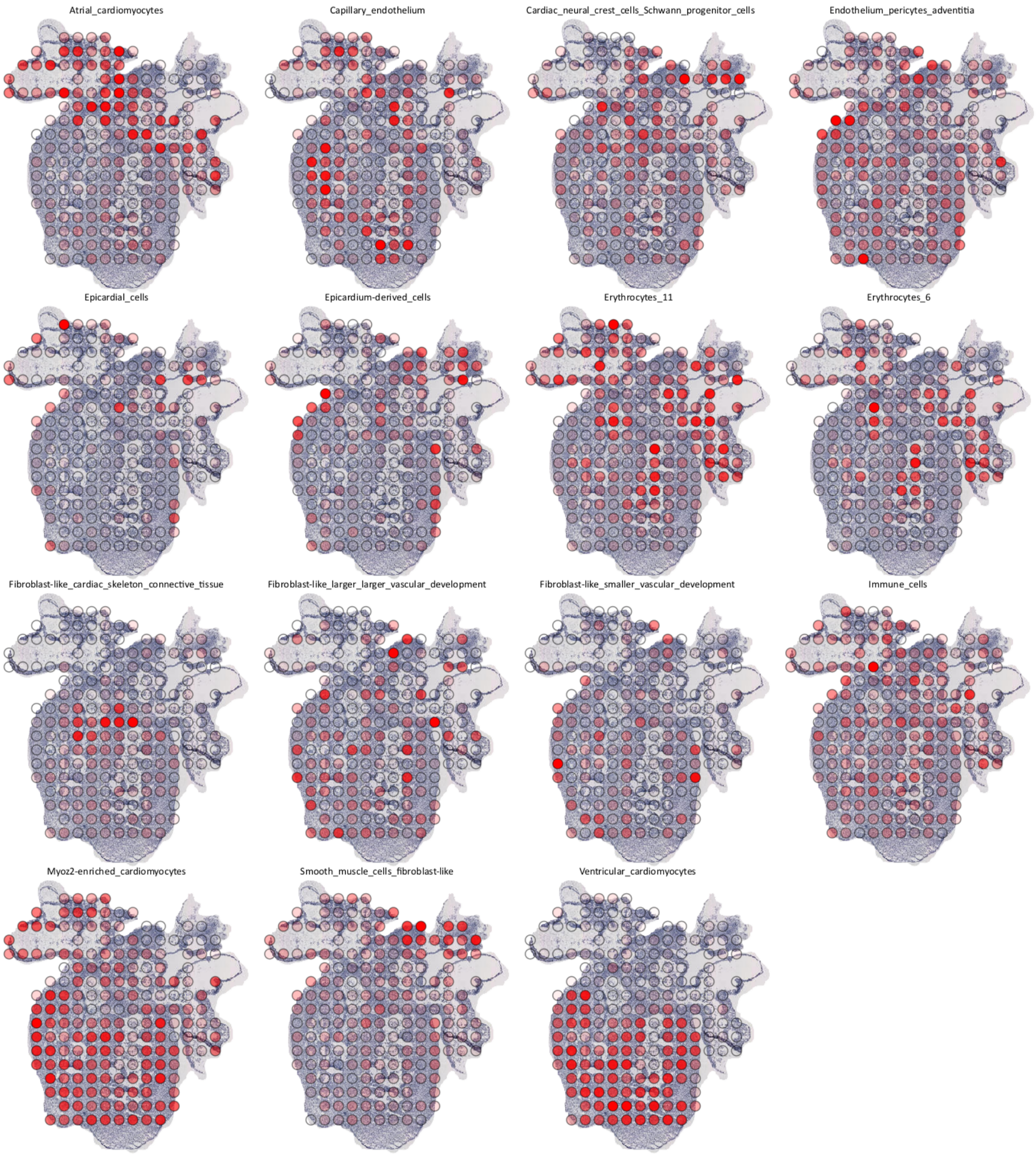


Supplementary Figure 1: Visualization of proportion estimates for section dh-A (from a series of eight independent sections from the same developmental heart, named A-H), scaled within each cell type.

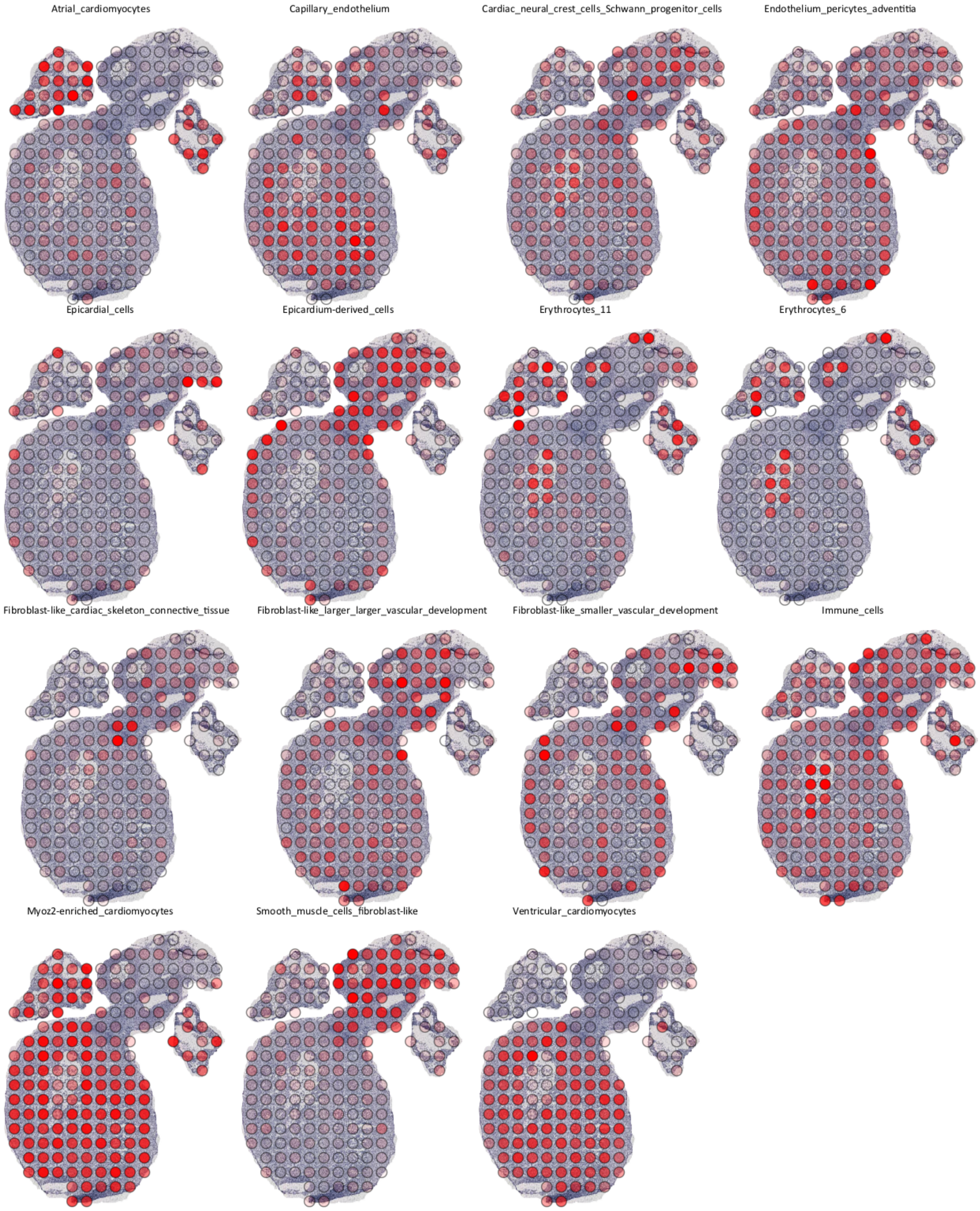


Supplementary Figure 2: Visualization of proportion estimates for section dh-B (from a series of eight independent sections from the same developmental heart, named A-H), scaled within each cell type.

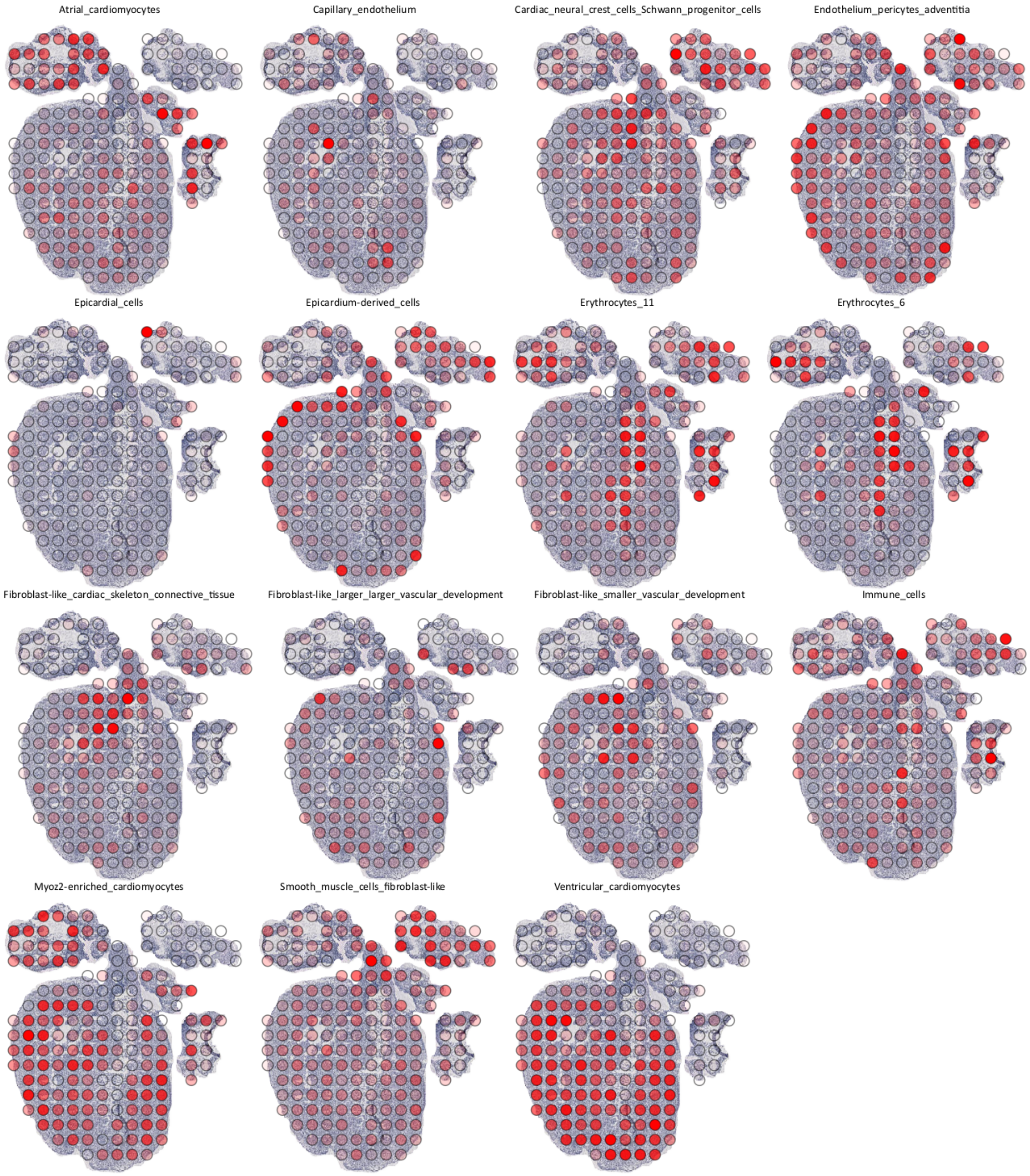


Supplementary Figure 3: Visualization of proportion estimates for section dh-C (from a series of eight independent sections from the same developmental heart, named A-H), scaled within each cell type.

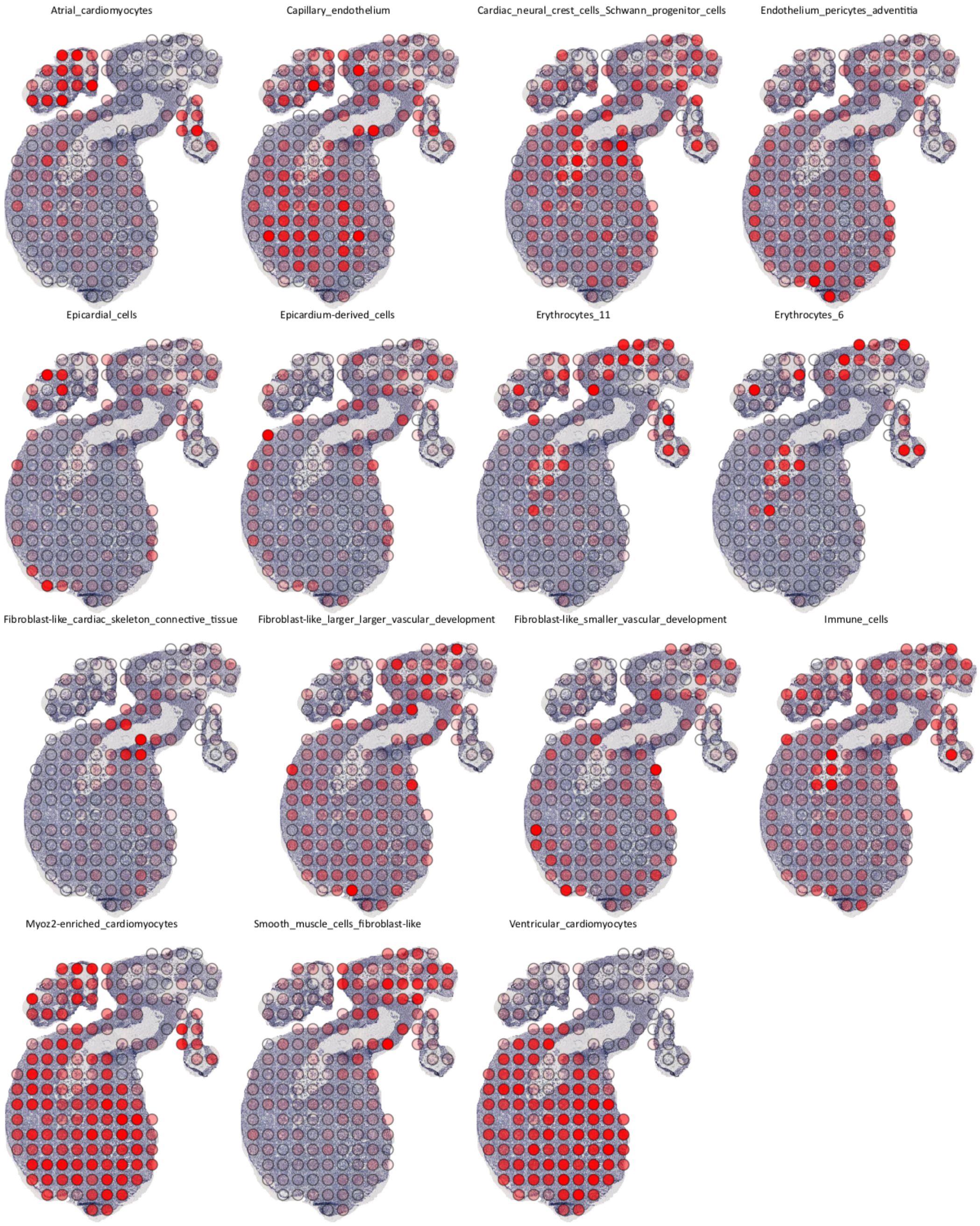


Supplementary Figure 4: Visualization of proportion estimates for section dh-D (from a series of eight independent sections from the same developmental heart, named A-H), scaled within each cell type.

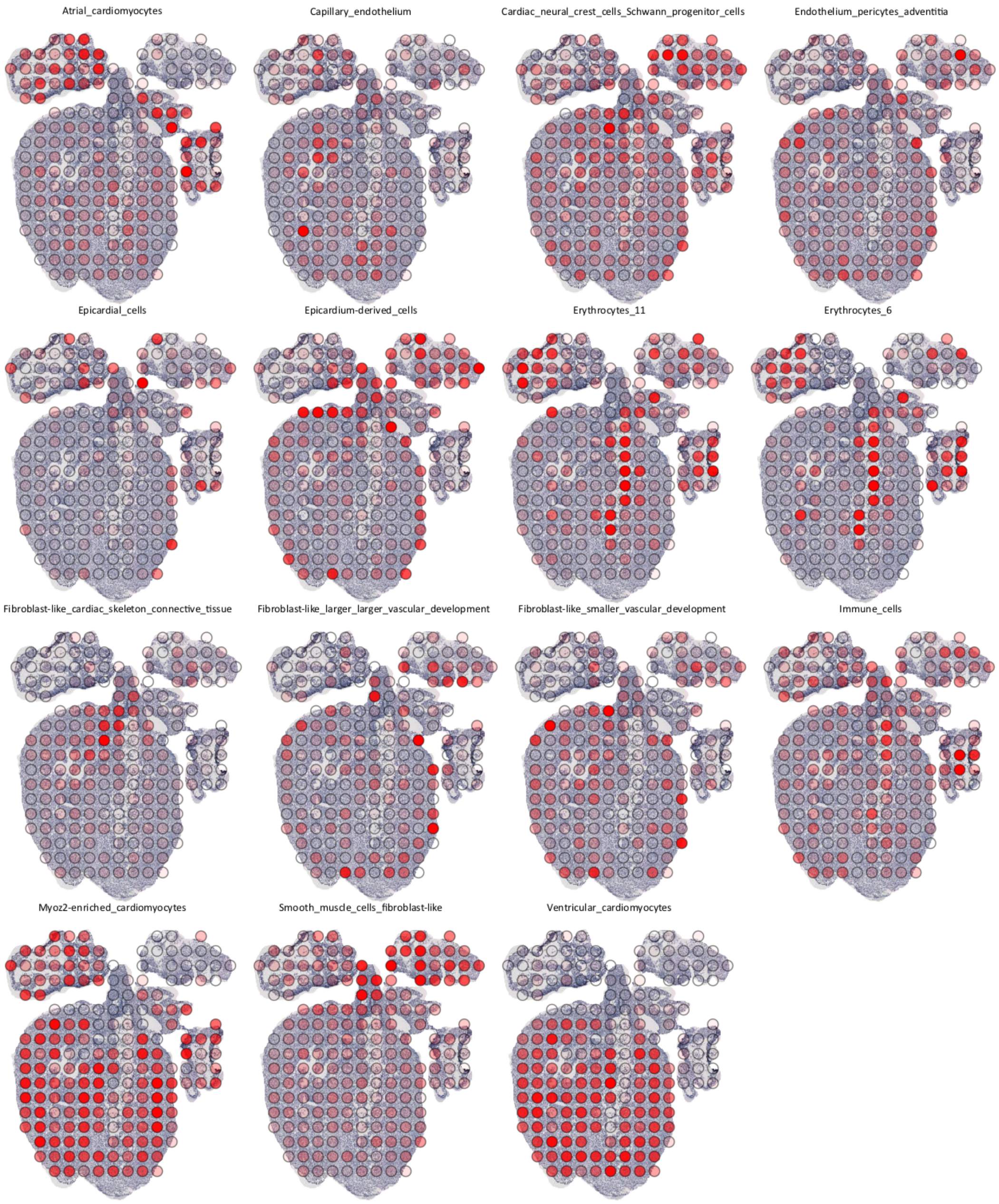


Supplementary Figure 5: Visualization of proportion estimates for section dh-E (from a series of eight independent sections from the same developmental heart, named A-H), scaled within each cell type.

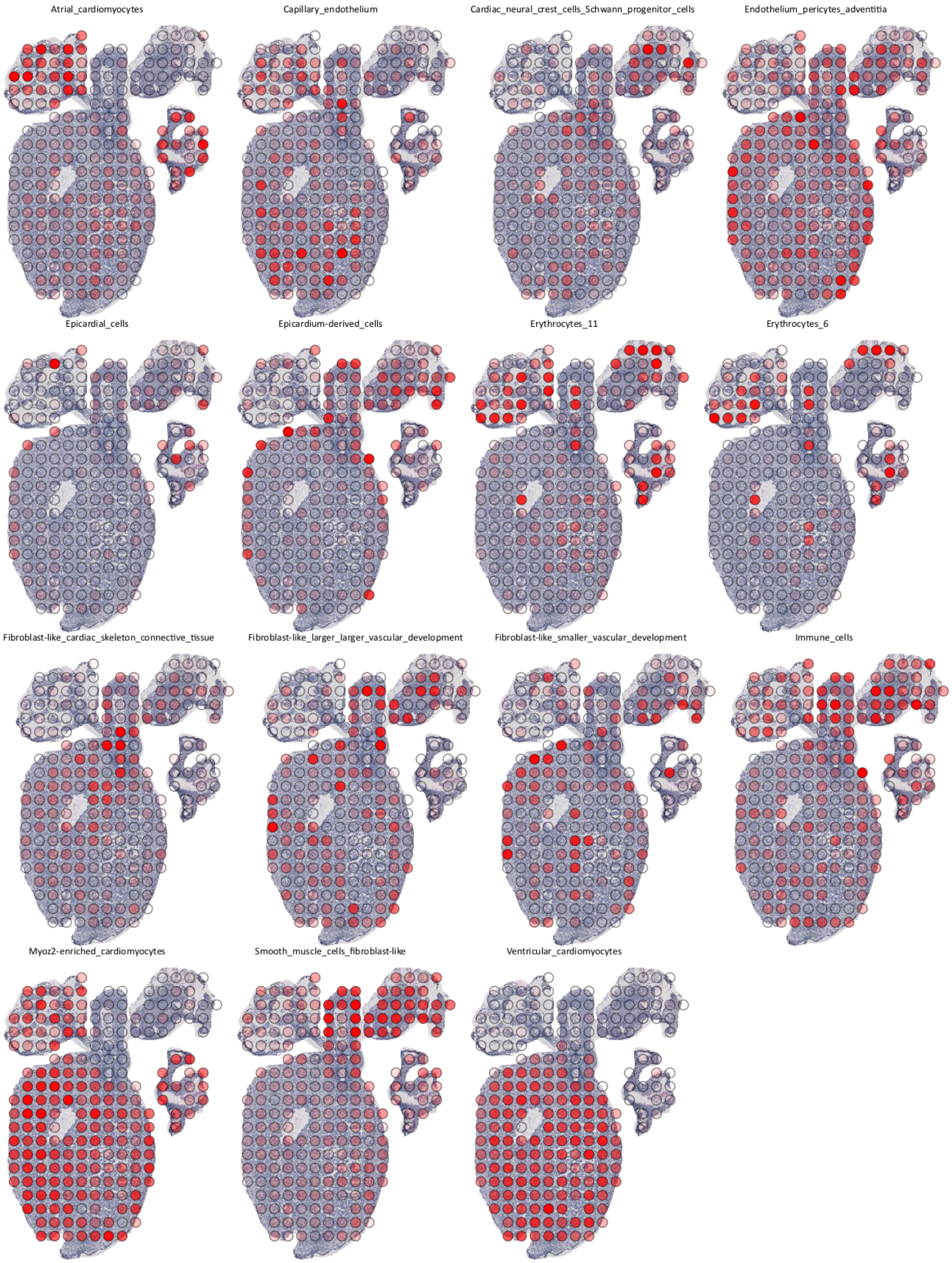


Supplementary Figure 6: Visualization of proportion estimates for section dh-F (from a series of eight independent sections from the same developmental heart, named A-H), scaled within each cell type.

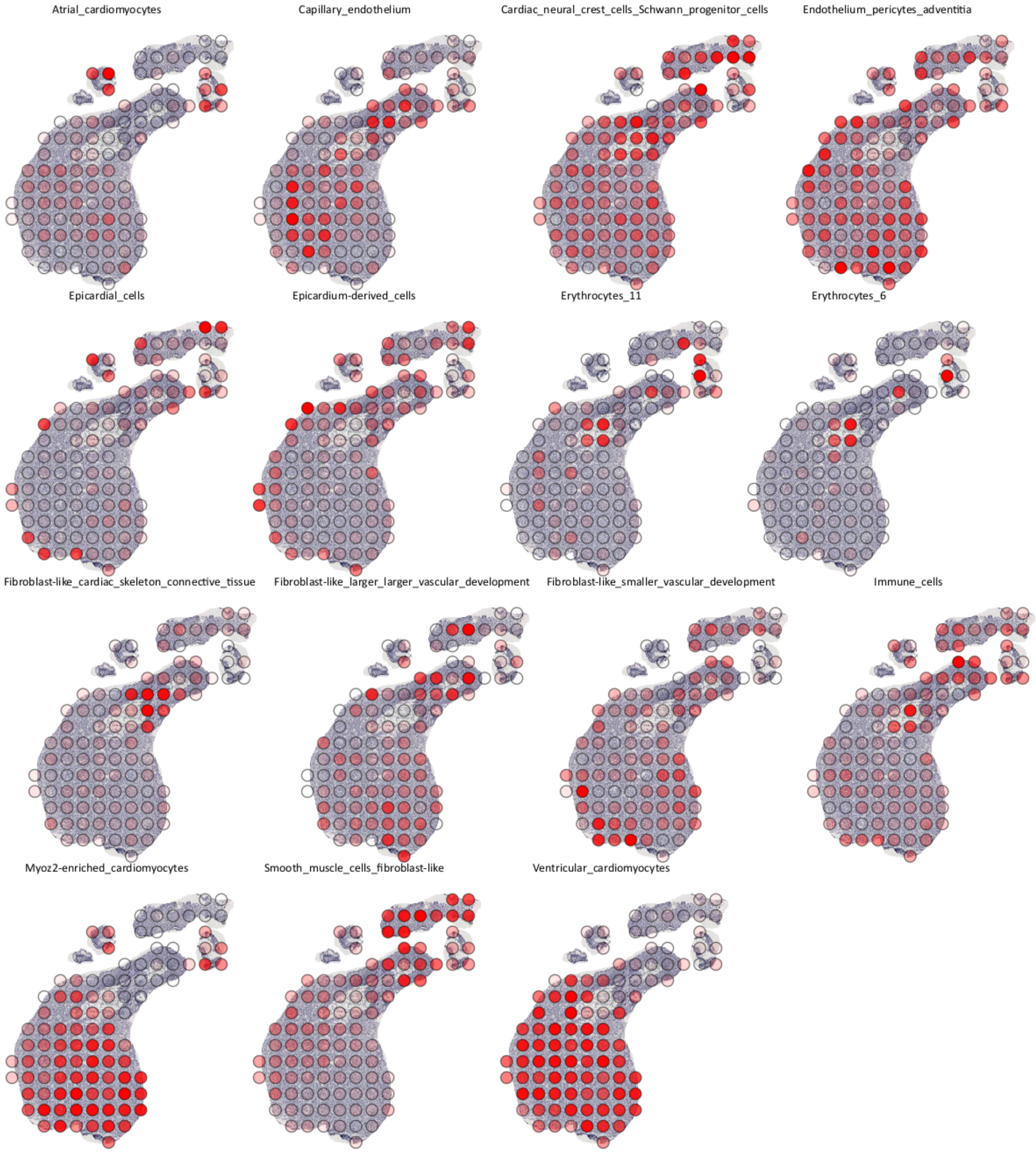


Supplementary Figure 7: Visualization of proportion estimates for section dh-G (from a series of eight independent sections from the same developmental heart, named A-H), scaled within each cell type.

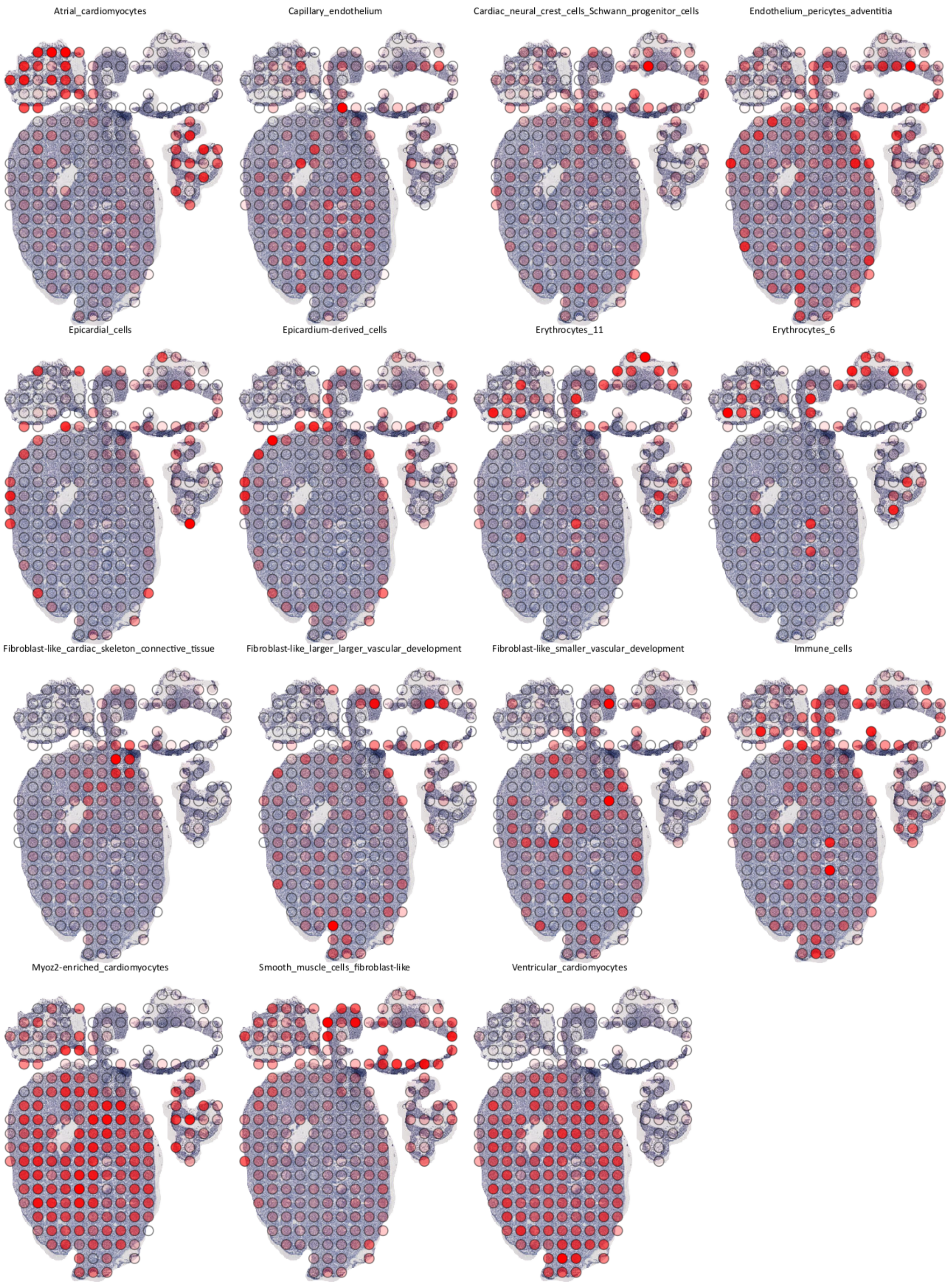


Supplementary Figure 8: Visualization of proportion estimates for section dh-H (from a series of eight independent sections from the same developmental heart, named A-H), scaled within each cell type.

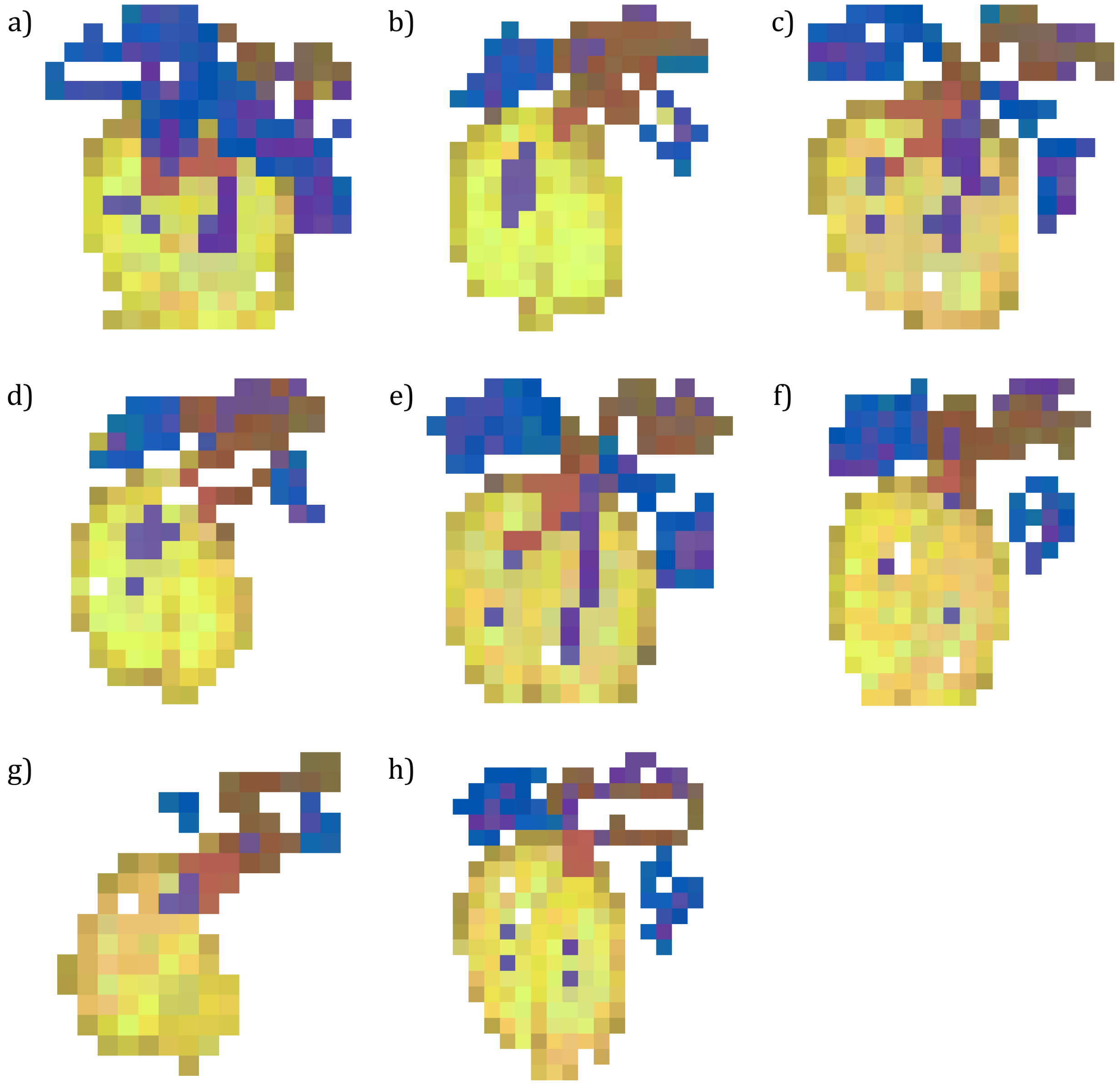


Supplementary Figure 9: Compressed cell type visualization of all sections of the developmental heart.

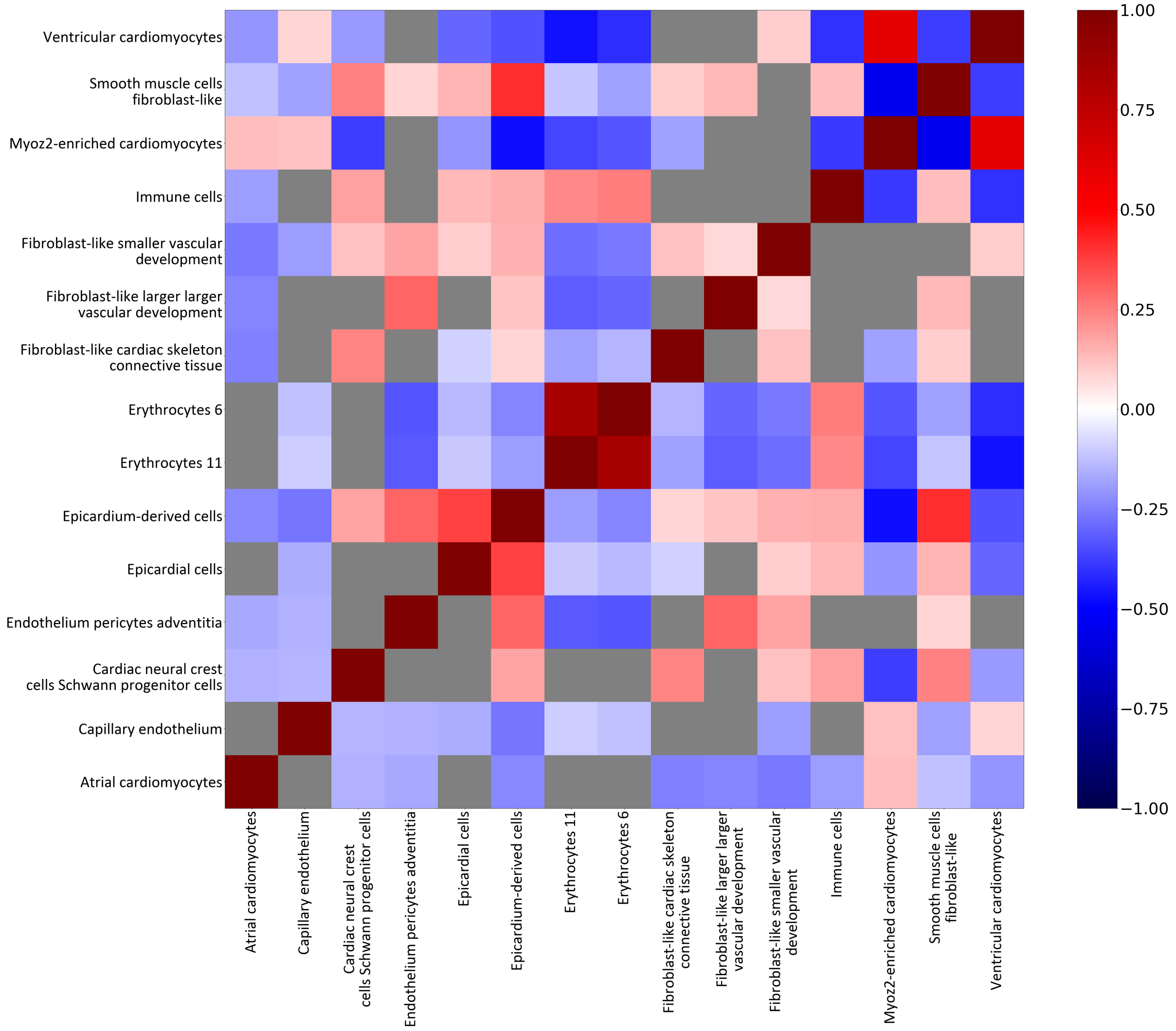


Supplementary Figure 10: Correlation between cell types (Methods) within the developmental heart. Gray areas represent correlation values which do not have a p-value below the significance threshold (0.01). The correlation values are computed over all 8 sections. Computed using the scipy function *scipy.stats.pearsonr*.

##### 1.2.2 Mouse Brain

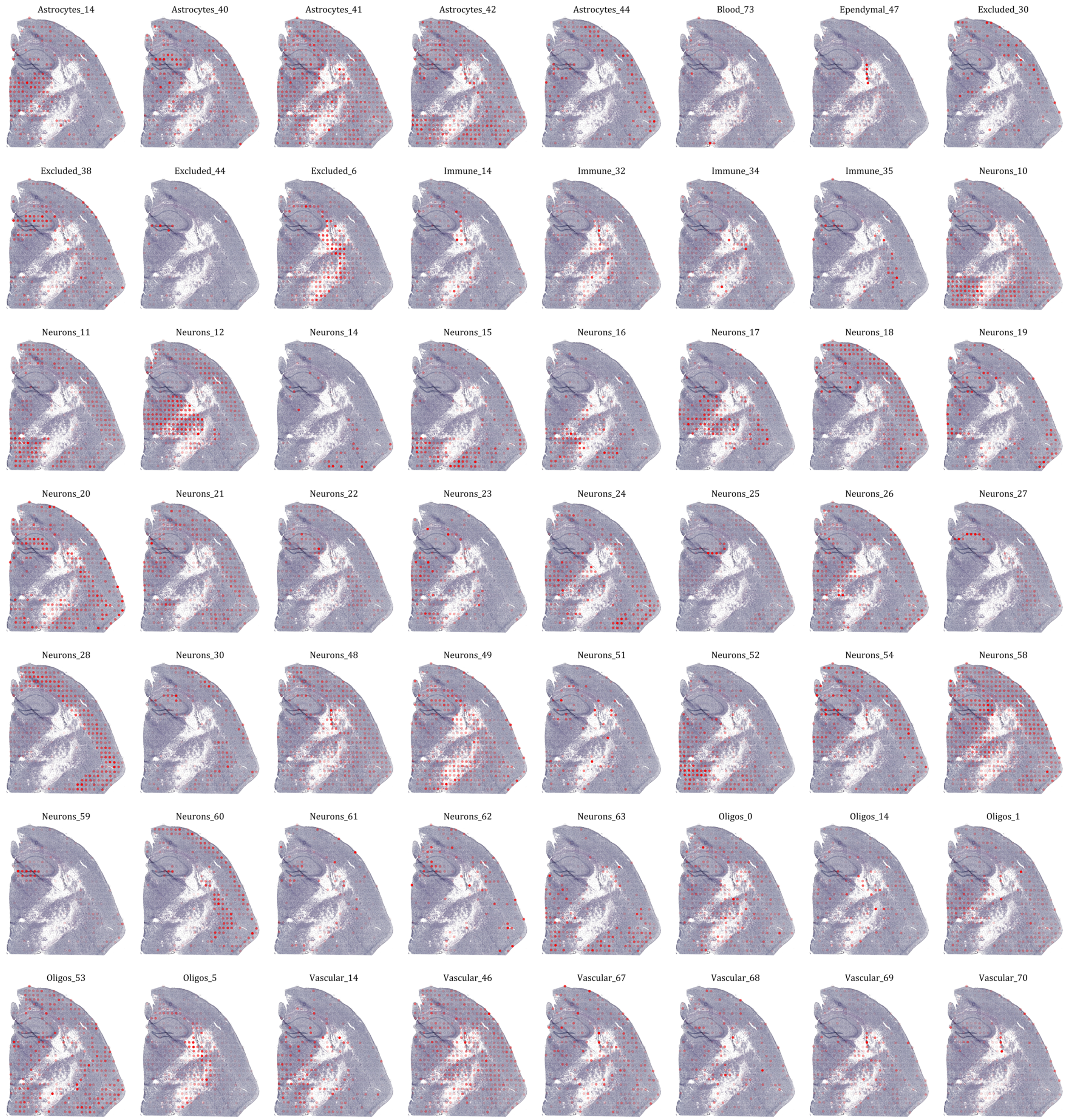


Supplementary Figure 11: Visualization of proportion estimates for section mb-A (ST array, 100 micron spots) of the mouse brain, scaled within each cell type.

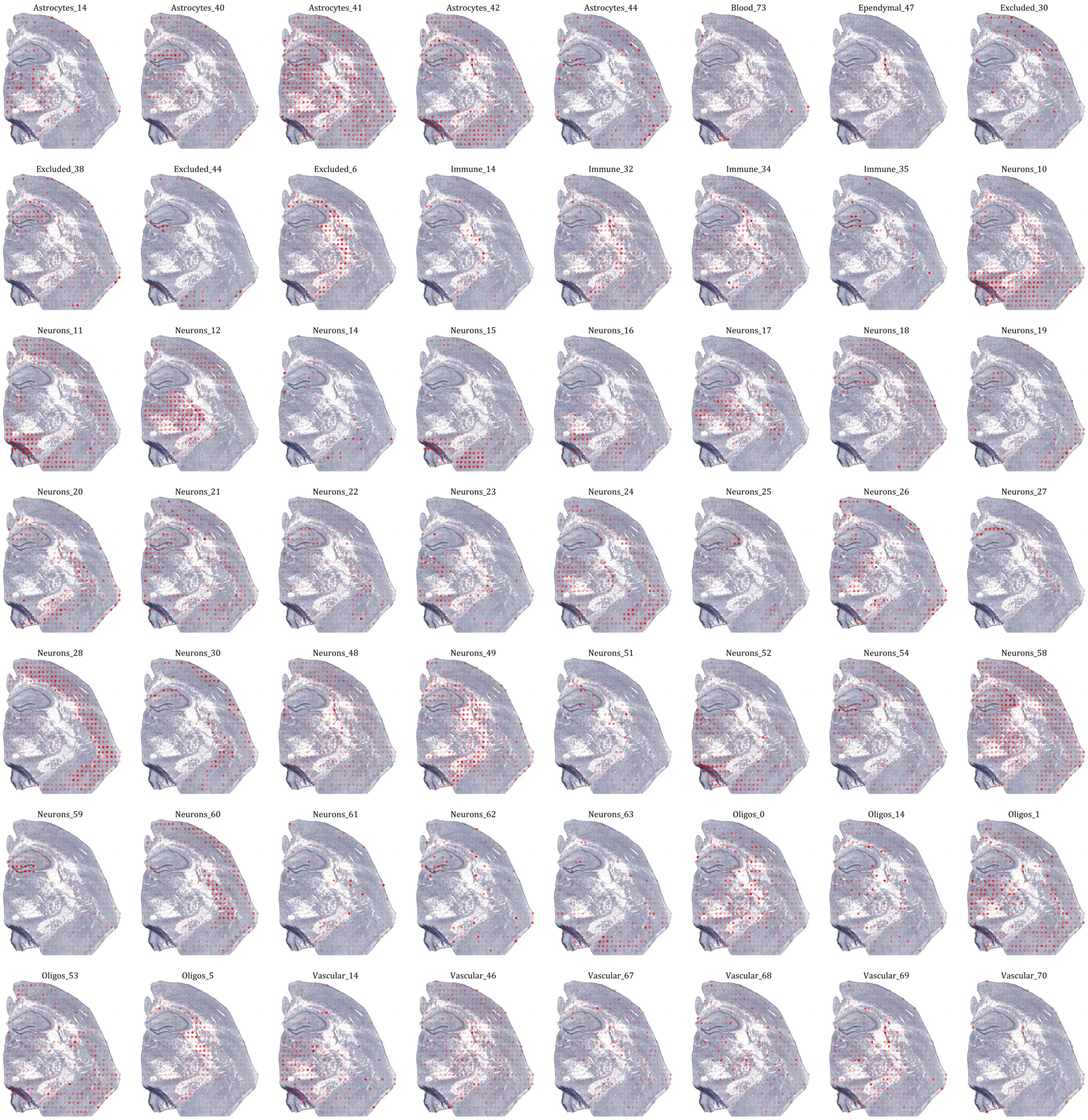


Supplementary Figure 12: Visualization of proportion estimates for section mb-$\alpha$ (ST array, 100 micron spots) of the mouse brain, scaled within each cell type.

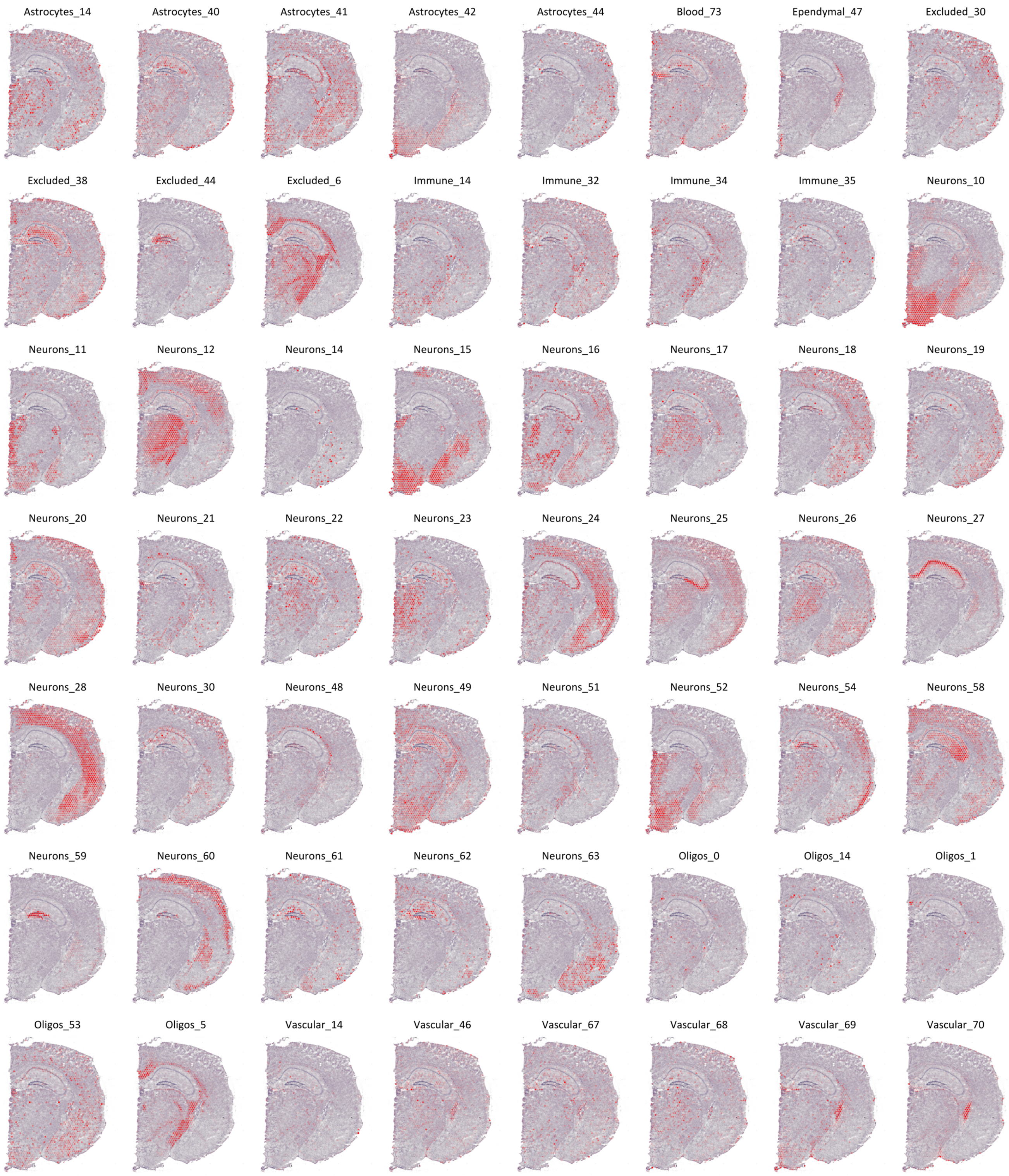


Supplementary Figure 13: Visualization of proportion estimates for section mb-B (Visium array, 55 micron spots), scaled within each cell type.

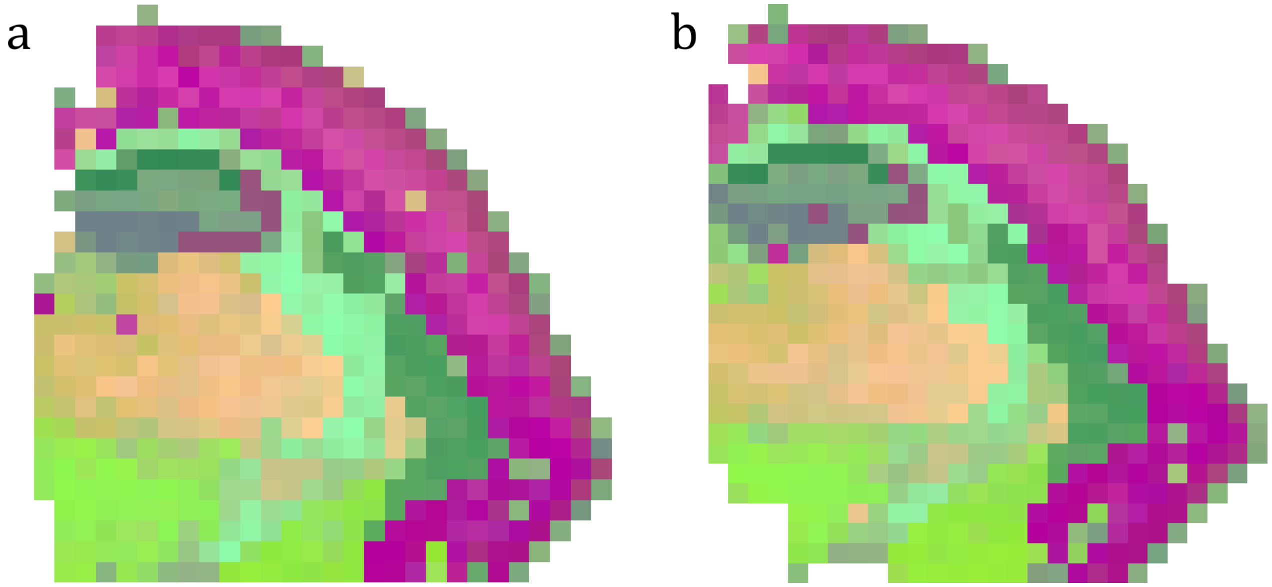


Supplementary Figure 14: Compressed cell type visualization of the mb-A (left) and mb-$\alpha$ (right) mouse brain sections (ST array, 100 micron spots).

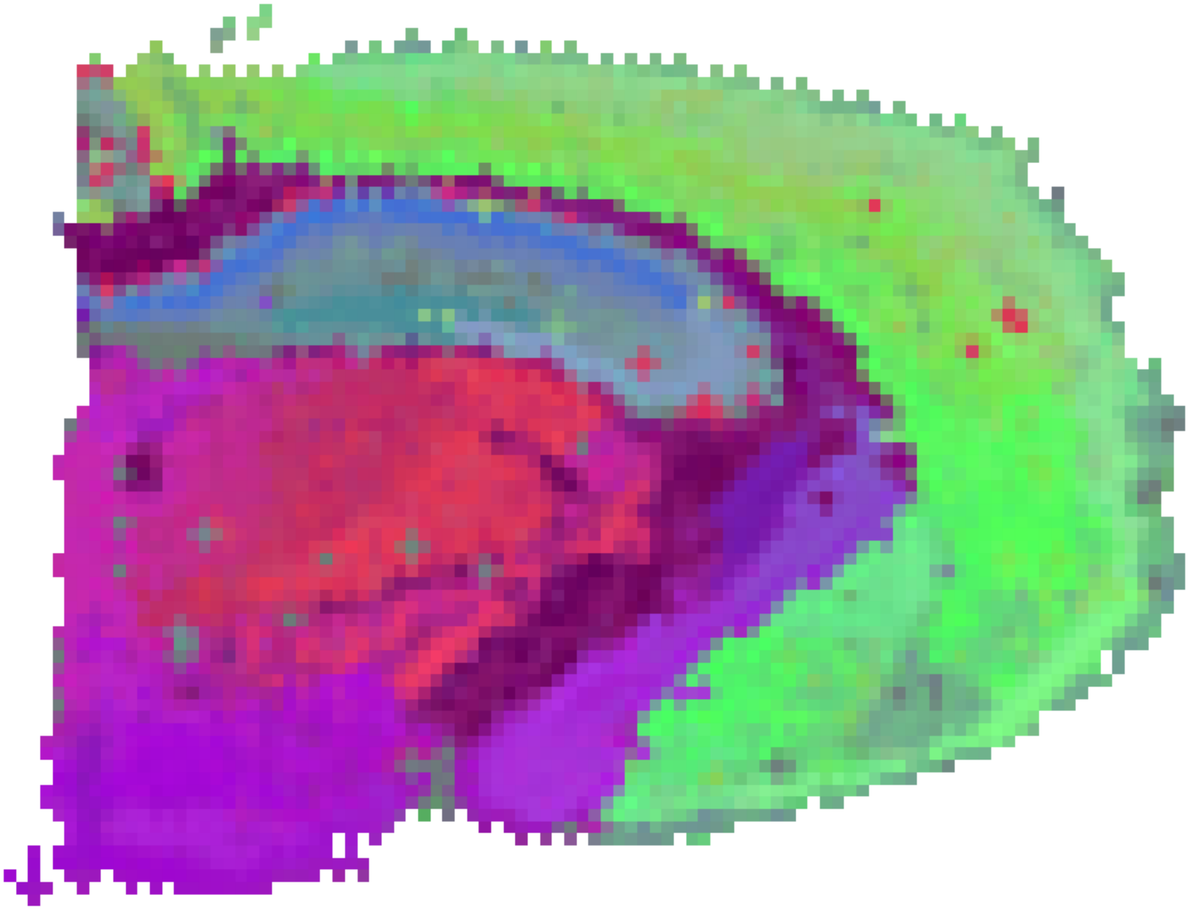

Supplementary Figure 15: Compressed cell type visualization of the mb-B (Visium array, 55 micron spots) mouse brain section.

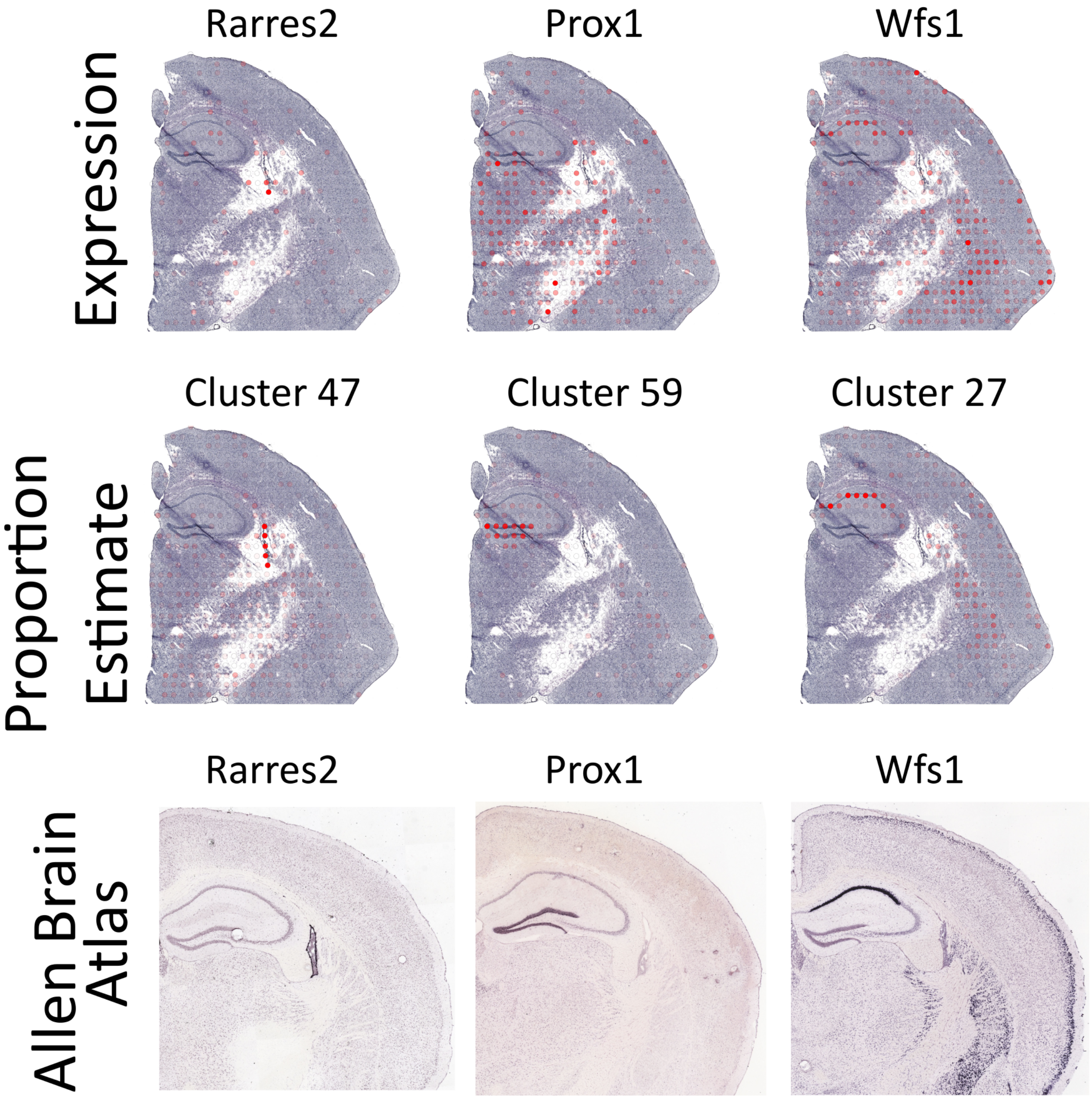


Supplementary Figure 16: Comparison between visualization of marker gene relative expression (top) and proportion estimate (middle) in section mb-A, with Allen Brain Atlas ISH images (bottom) as reference. The relative gene expression is obtained by dividing the number of observed transcripts at a given spot ($x_{sg}$) by the total number of observed transcripts in the given spot. The relative gene expression values are visualized according to the same procedure as the proportion values (Methods).

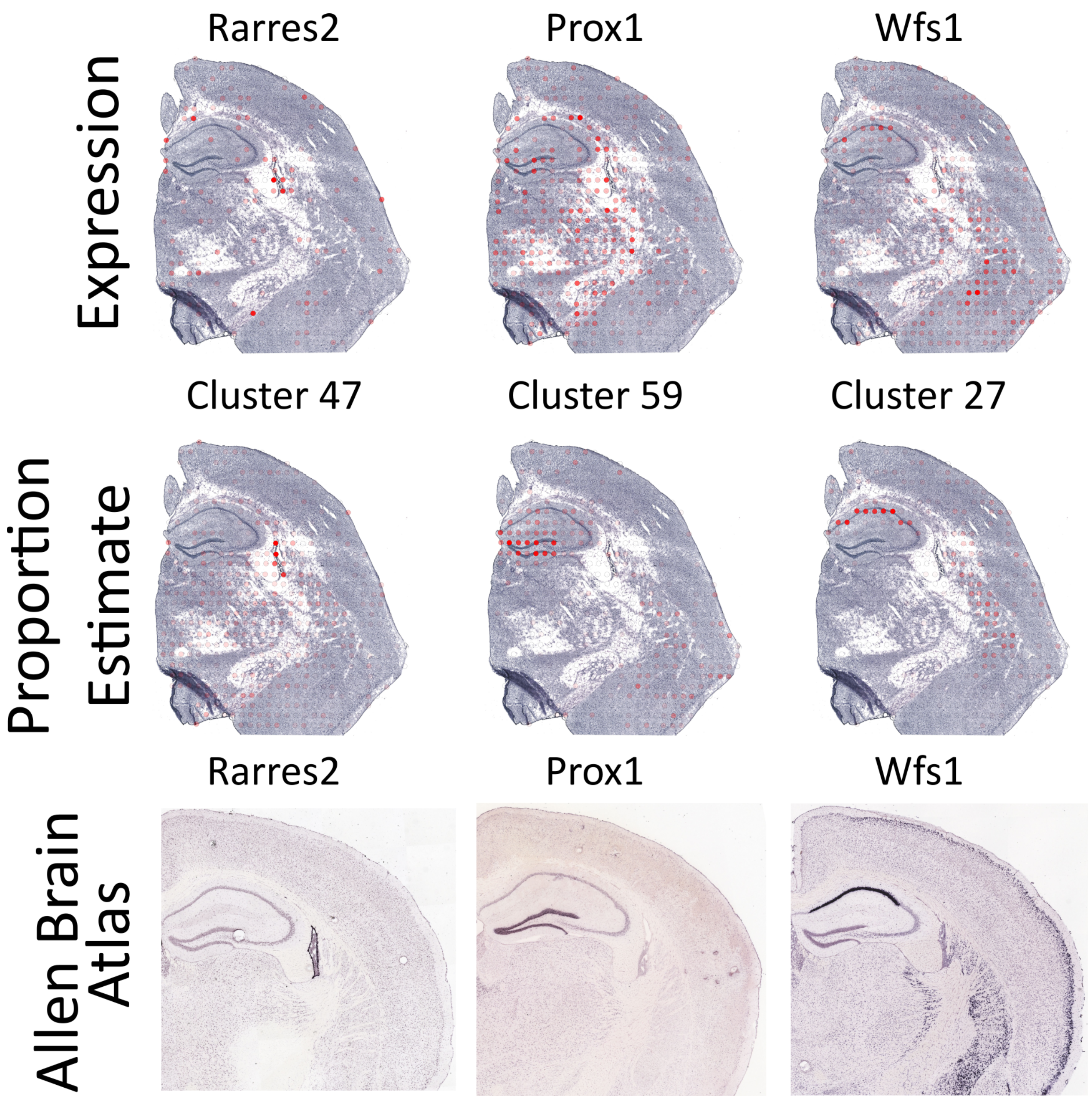


Supplementary Figure 17: Comparison between visualization of marker gene relative expression (top) and proportion estimate (middle) in section mb-$\alpha$, with Allen Brain Atlas ISH images (bottom) as reference. The relative gene expression is obtained by dividing the number of observed transcripts at a given spot ($x_{sg}$) by the total number of observed transcripts in the given spot. The relative gene expression values are visualized according to the same procedure as the proportion values (Methods).

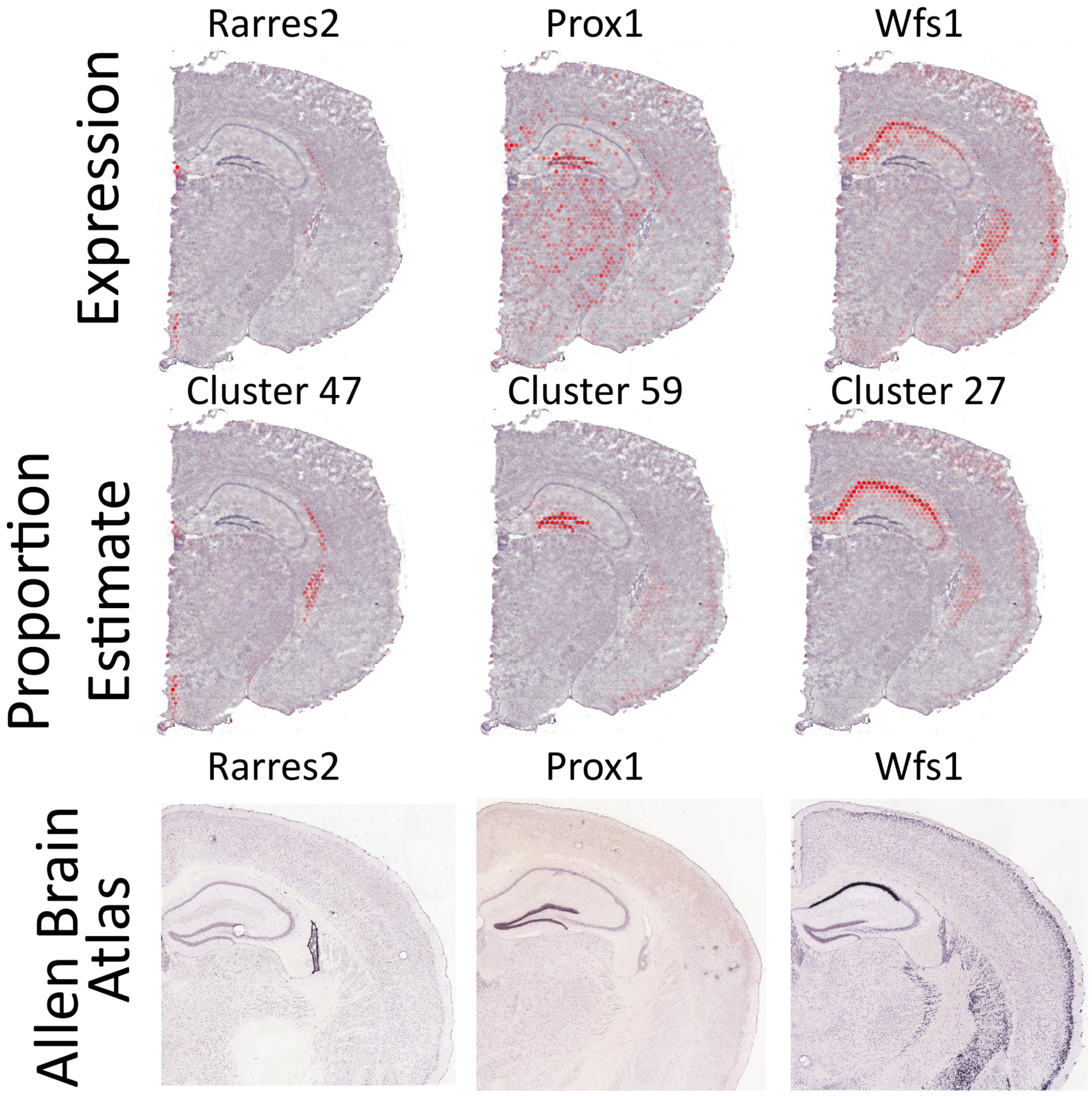


Supplementary Figure 18: Comparison between visualization of marker gene relative expression (top) and proportion estimate (middle) in section mb-B, with Allen Brain Atlas ISH images (bottom) as reference. The relative gene expression is obtained by dividing the number of observed transcripts at a given spot ($x_{sg}$) by the total number of observed transcripts in the given spot. The relative gene expression values are visualized according to the same procedure as the proportion values (Methods).

##### 1.2.3 Comparison

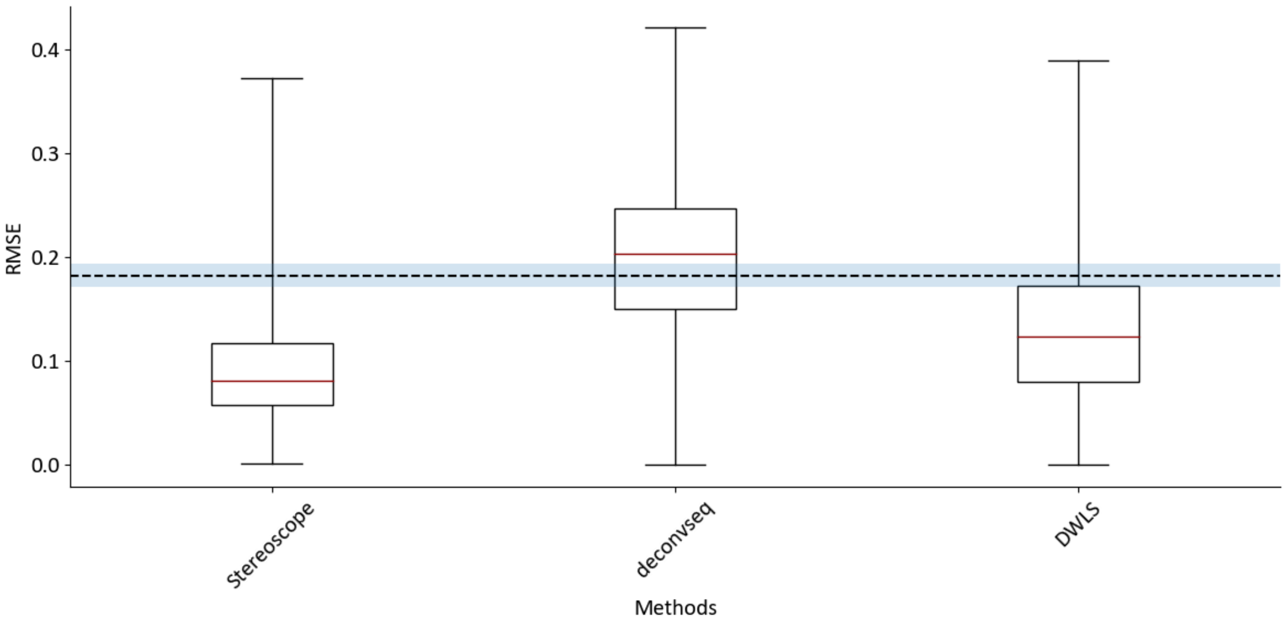


Supplementary Figure 19: Comparison between Stereoscope and other methods designed to estimate proportions of cell types in bulk data with the help of single cell data. The red line indicates the median and the whiskers show the full range of the values. The dashed line indicates the average value from computing the RMSE from 1000 proportion estimates (of each spot) generated from a Dirichlet distribution (concentration set to 1 for all types). The shaded blue regions show the 95% confidence intervals of the mean value.

| **Compared to** | **Mean Difference** | **p-value** |
|---|---|---|
| deconvseq | -0.102095 | 6.562620e-107 |
| DWLS | -0.039784 | 2.966613e-77 |

Supplementary Table 4: Results from a one-sided Wilcoxon signed rank test – effectively testing whether the difference between the spotwise paired RMSE values are symmetrically distributed around zero or if this distribution is skewed in favor of Stereoscope. The mean difference is computed as the average difference between Stereoscope and the other method's RMSE values in each spot, where a negative value means that Stereoscope on average have lower RMSE values.
